# A comparative genomic analysis of the barley pathogen *Pyrenophora teres* f. *teres* identifies sub-telomeric regions as drivers of virulence

**DOI:** 10.1101/753202

**Authors:** Nathan A. Wyatt, Jonathan K. Richards, Robert S. Brueggeman, Timothy L. Friesen

## Abstract

*Pyrenophora teres* f. *teres* causes net form net blotch of barley and is an economically important pathogen throughout the world. However, *P. teres* f. *teres* is lacking in the genomic resources necessary to characterize the mechanisms of virulence. Recently a high quality reference genome was generated for *P. teres* f. *teres* isolate 0-1. Here, we present the reference quality sequence and annotation of four new isolates and we use the five available *P. teres* f. *teres* genomes for an in-depth comparison resulting in the generation of hypotheses pertaining to the potential mechanisms and evolution of virulence. Comparative analyses were performed between all five *P. teres* f. *teres* genomes examining genomic organization, structural variations, and core and accessory genomic content, specifically focusing on the genomic characterization of known virulence loci and the localization of genes predicted to encode secreted and effector proteins. We showed that 14 of 15 currently published virulence quantitative trait loci (QTL) span accessory genomic regions consistent with these accessory regions being important drivers of host adaptation. Additionally, these accessory genomic regions were frequently found in sub-telomeric regions of chromosomes with 10 of the 14 accessory region QTL localizing to sub-telomeric regions. Comparative analysis of the sub-telomeric regions of *P. teres* f. *teres* chromosomes revealed translocation events where homology was detected between non-homologous chromosomes at a significantly higher rate than the rest of the genome. These results indicate that the sub-telomeric accessory genomic compartments not only harbor most of the known virulence loci, but also that these regions have the capacity to rapidly evolve.

## Background

The intimate process of co-evolution between plants and their pathogens is often depicted as a form of trench warfare where pathogens evolve to manipulate the host, leading to the acquisition of nutrients and in response, selection pressure is placed on the plant to develop mechanisms to resist the pathogen (Stahl and Bishop 2000). Plants have evolved an innate immune system that recognizes and responds to microbes that successful pathogens must overcome (Cook et al. 2015; Jones and Dangle 2006). This physical interaction of a pathogen with its host often occurs through a set of small secreted proteins termed “effectors”.

Pathogen effectors have evolved to manipulate the host defense response to the benefit of the pathogen, but these effectors also provide targets for plant recognition of the pathogen (Bialas et al. 2017). A pathogen’s effector function is specific to the pathogen’s lifestyle and mode of nutrient acquisition. Obligate biotrophic pathogens typically invade host tissue in a manner designed to evade host recognition to gain nutrient from living tissue. Obligate biotrophs often develop haustoria that are appendages of fungal hyphae used to invaginate plant cells to serve as feeding structures and effector secretion hot spots (Perfect et al. 2001). Conversely, other pathogens, often classified as necrotrophs, can gain nutrient from dead or dying tissue resulting from pathogen induced programmed cell death of plant cells and have evolved mechanisms to deal with the various aspects of the plant defense response (Liu et al. 2012b; Liu et al. 2016). Between these two models is a spectrum of fungi often classified as hembiotrophic that begin plant colonization in a manner similar to a biotrophic or endophytic interaction, referred to as the biotrophic or symptomless phase, where evasion of plant recognition is important. The symptomless or biotrophic phase is followed by a transition to the necrotrophic phase, characterized by inducing host cell death, colonization, and eventual sporulation (Vleeshouwers et al. 2014).

Effector discovery is hindered by a lack of homology to known proteins and the association of genes that encode effectors with low complexity repeat-rich genomic compartments that are drivers of rapid evolutionary adaptation (Raffaele and Kamoun 2012; Dong et al. 2015; Faino et al. 2016). Pathogens must be able to quickly modify or lose effectors that are co-opted by the host, resulting in effector genes that can be highly polymorphic between members of the same species (Cook et al. 2015). Low complexity regions, that are rich in repeats and harbor transposable elements (TEs), are thought to be dynamic compartments involved in rapid evolution as compared to the more conserved stable gene-dense compartments. Pathogen effector genes often reside in these low complexity regions and consequently are affected by increased structural polymorphism, point mutagenesis, and diversifying selection that are common to these genomic compartments (Raffaele 2010; Rouxel 2011; de Jonge et al. 2013; Dong et al. 2015; Faino et al. 2016, Moller and Stukenbrock 2017).

Generally, in fungi, the lack of available sequence data has been attributed to the difficulties and costs of sequencing and assembling genomes (Thomma et al. 2016). The Pacific Biosciences (PacBio) single molecule real-time (SMRT) sequencing platform is capable of read lengths of up to 60 Kb allowing for the sequencing and assembly of full microbial genomes containing large repetitive elements (Goodwin et al. 2016; Thomma et al. 2016). In recent years, SMRT sequencing has been used to obtain complete genome sequences of a number of fungal plant pathogen species, demonstrating its utility for fungal plant pathogen genomics (Faino et al. 2015; Thomma et al. 2016; van Kan et al. 2016, Derbyshire et al. 2017; Wyatt et al. 2018; Richards et al. 2018). Finishing fungal genomes opens the door to identifying all of the genes in the genome including previously difficult to identify effector genes residing in low complexity regions. When multiple isolates of the same species can be fully sequenced, the full gene repertoires of the species can be compared and characterized, this is termed the pan-genome. (Hurgobin & Edwards 2017; Tettelin et al. 2005, Vernikos et al. 2015). This pan-genome defines the entire genomic repertoire of a given taxonomic clade and accounts for all possible lifestyle processes of the organisms being observed. Pan-genomes are further defined by their core genomes, representing genes common to all individuals, and their accessory genomes, representing genes that are unique to individuals within the clade. Core genomes often encode proteins associated with basic biological aspects of the entire clade, whereas accessory genomes are thought to encode genes involved in niche adaptation. The intra-specific gene content of the accessory genome is of particular interest to plant-microbe interactions given that the genes responsible for virulence are often not shared among all individuals of a species (Croll et al. 2012). With the decreased cost of sequencing technology, it has become possible to sequence many individuals of a species to compare intraspecific gene content and structural variation. These comparative studies have been done for a number of bacterial species as well as some well-studied crop species. Most recently, pan-genome analyses have been conducted in the fungal plant pathogen species *Zymoseptoria tritici* (Plissonneau et al. 2018)*, Pyrenophora tritici-repentis* (Moolhuijzen et al. 2018), and *Parastagonospora nodorum* (Syme et al. 2018a).

Net form net blotch (NFNB) is a stubble born foliar disease of barley (*Hordeum vulgare*) induced by the fungal pathogen *Pyrenophora teres* f. *teres*. Typical disease losses due to NFNB have ranged between 10 and 40% with the potential for complete yield loss given environmental conditions favorable to the pathogen, namely, wide planting of a susceptible cultivar, and high humidity or rainfall (Mathre 1997). *P*. *teres* f. *teres* infection on a susceptible host results in dot-like lesions on the leaf, progressing to longitudinal striations and finally the net-like pattern from which the disease gets its name. Necrotrophic effectors (NEs) were shown to be an important component of the *P*. *teres* f. *teres* infection cycle (Liu et al. 2015), similar to the closely related species *P*. *nodorum* and *P*. *tritici-repentis* that also employ NEs to induce NE triggered susceptibility (NETS) (Faris et al. 2013; Friesen and Faris 2010). In addition to NETS, dominant resistance has also been identified in a number of barley backgrounds that follow a gene-for-gene model. The presence of both dominant resistance and dominant susceptibility and the discovery of NEs illustrates the complicated interaction at play (Liu et al. 2011; Liu et al. 2012a; Liu et al. 2015; Koladia et al. 2017). As of yet, no effector genes have been cloned and characterized in *P*. *teres* f. *teres* and therefore the mechanism of virulence/pathogenicity is still poorly understood.

Adding to the complexity of *P*. *teres* f. *teres* virulence is the diversity observed in both local and global pathogen populations. Host genotype specificity was first observed in a set of Australian *P*. *teres* f. *teres* isolates (Khan and Boyd 1969) and bi-parental pathogen mapping populations have been an important tool used to associate genomic loci of *P*. *teres* f. *teres* isolates with host specific virulence and avirulence (Weiland et al. *1999*; Lai et al. 2007; Beattie et al. 2007; Liu et al. 2011; Shjerve et al. 2014; Koladia et al. 2017). Currently published mapping populations have been developed from crosses of *P. teres* f. *teres* isolates 0-1 × 15A, isolates 15A × 6A, and isolates FGOH04Ptt-21 × BB25 and report on a total of 15 unique genetic loci contributing to virulence leading to NFNB disease on different barley lines.

The diversity observed in *P*. *teres* f. *teres* bi-parental mapping studies presents an obstacle for effector discovery in *P*. *teres* f. *teres* isolates that have virulence associations not present in the currently published reference isolate 0-1 (Ellwood et al. 2010; Wyatt et al. 2018). With each new bi-parental mapping study published, unique QTL have been identified on common cultivars located in different parts of the *P. teres* f. *teres* genome with very few QTL being identified in all bi-parental mapping studies (Weiland *et al*. 1999; Lai *et al*. 2007; Shjerve *et al*. 2014; Koladia *et al*. 2017). Given the diversity observed in previous mapping studies of avirulence/virulence, it is unlikely that a single isolate would capture a representative sample of the effectorome of *P*. *teres* f. *teres*. To examine a broader sampling of the *P*. *teres* f. *teres* effectorome, we used a pan-genome approach by sequencing and assembling four additional isolates using PacBio SMRT sequencing and annotated their genomes using RNAseq support following the same protocol outlined in Wyatt et al. (2018). Isolates sequenced have all been used previously in biparental mapping studies including the two California, USA, isolates 15A (Weiland et al. 1999; Lai et al. 2007; Shjerve et al. 2014) and 6A (Shjerve et al. 2014); the North Dakota, USA, isolate FGOH04Ptt-21 (Koladia et al. 2017), and the Danish isolate BB25 (Koladia et al. 2017). Using the reference isolate 0-1 (Lai et al. 2007; Ellwood et al. 2010; Wyatt et al. 2018) and the four newly sequenced isolates, a pan genome analyses was performed on the five isolates to evaluate the diversity in genomic architecture and gene content. The genome annotations produced were used to identify chromosome structural variation and parse the effectorome of each isolate for comparison of effector diversity within *P*. *teres* f. *teres* isolates to examine common and unique effectors of the species.

## Results

### Genome sequencing, assembly, and scaffolding

To assess genome composition and structure of *P. teres* f. *teres*, we generated high quality genome assemblies using PacBio SMRT sequencing for isolates 15A, 6A, FGOH04Ptt-21, and BB25. These isolates were chosen due to their differing disease reactions on commonly used differential barley lines and their inclusion in previously reported genetic studies using pathogen bi-parental mapping populations (Weiland et al. 1999; Lai et al. 2007; Shjerve et al. 2014; Koladia et al. 2017). Sequencing results of the four new *P. teres* f. *teres* isolates compare favorably with previous sequencing efforts for *P*. *teres* f. *teres* isolate 0-1 which produced 1,148,507 reads with average read lengths of 8,051 bps (Wyatt et al. 2018) (Table 1).

**Table 1.**
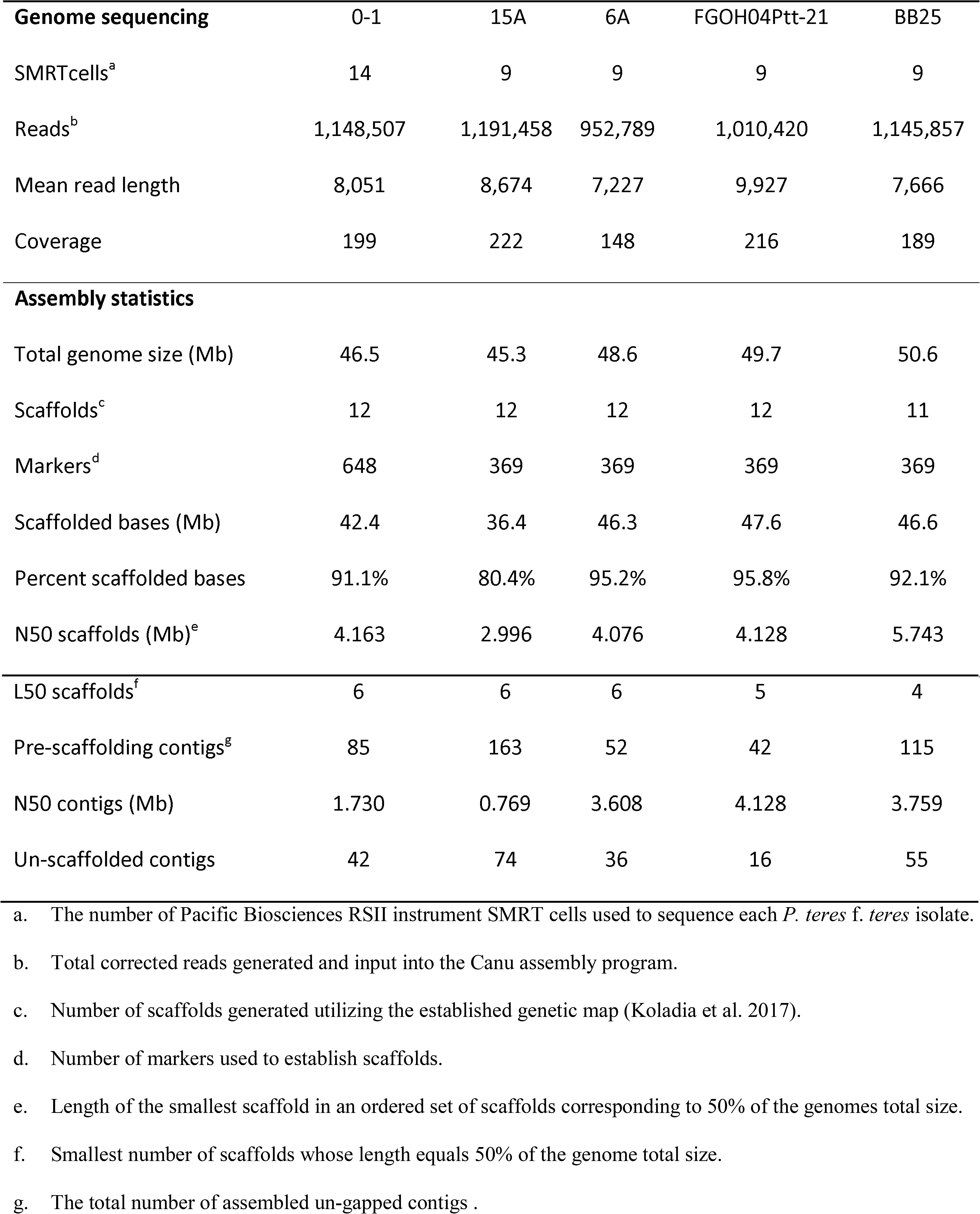
Summary statistics for P. teres f. teres genome assemblies.

| **Genome sequencing** | 0-1 | 15A | 6A | FGOH04Ptt-21 | BB25 |
| --- | --- | --- | --- | --- | --- |
| SMRTcells<sup>a</sup> | 14 | 9 | 9 | 9 | 9 |
| Reads<sup>b</sup> | 1,148,507 | 1,191,458 | 952,789 | 1,010,420 | 1,145,857 |
| Mean read length | 8,051 | 8,674 | 7,227 | 9,927 | 7,666 |
| Coverage | 199 | 222 | 148 | 216 | 189 |
| **Assembly statistics** |  |  |  |  |  |
| Total genome size (Mb) | 46.5 | 45.3 | 48.6 | 49.7 | 50.6 |
| Scaffolds<sup>c</sup> | 12 | 12 | 12 | 12 | 11 |
| Markers<sup>d</sup> | 648 | 369 | 369 | 369 | 369 |
| Scaffolded bases (Mb) | 42.4 | 36.4 | 46.3 | 47.6 | 46.6 |
| Percent scaffolded bases | 91.1% | 80.4% | 95.2% | 95.8% | 92.1% |
| N50 scaffolds (Mb)<sup>e</sup> | 4.163 | 2.996 | 4.076 | 4.128 | 5.743 |
| L50 scaffolds<sup>f</sup> | 6 | 6 | 6 | 5 | 4 |
| Pre-scaffolding contigs<sup>g</sup> | 85 | 163 | 52 | 42 | 115 |
| N50 contigs (Mb) | 1.730 | 0.769 | 3.608 | 4.128 | 3.759 |
| Un-scaffolded contigs | 42 | 74 | 36 | 16 | 55 |

a. The number of Pacific Biosciences RSII instrument SMRT cells used to sequence each *P. teres f. teres* isolate.
b. Total corrected reads generated and input into the Canu assembly program.
c. Number of scaffolds generated utilizing the established genetic map (Koladia et al. 2017).
d. Number of markers used to establish scaffolds.
e. Length of the smallest scaffold in an ordered set of scaffolds corresponding to 50% of the genomes total size.
f. Smallest number of scaffolds whose length equals 50% of the genome total size.
g. The total number of assembled un-gapped contigs .

Assemblies of the sequenced *P*. *teres* f. *teres* isolates 15A, 6A, FGOH04Ptt-21, and BB25 also compared favorably with the currently published *P*. *teres* f. *teres* isolate 0-1 assembly having assembled into 163, 52, 42, and 112 contigs, respectively (Table1). Scaffolding of the resulting assemblies using the 16 linkage groups of the FGOH04Ptt-21 × BB25 linkage maps (Koladia et al. 2017) produced 12 total scaffolds for *P*. *teres* f. *teres* isolates 15A, 6A, and FGOH04Ptt-21 and produced 11 scaffolds for *P*. *teres* f. *teres* isolate BB25 (Table1). This is a reduced number of scaffolds compared to the number of linkage groups due to assembled genomic sequence coverage across multiple linkage groups. *P. teres* f. *teres* has previously been reported to have 12 chromosomes and the 12 and 11 scaffolds in the four newly sequenced isolates represent near full chromosomes based on alignments to previously assembled *P. teres* f. *teres* genomes (Wyatt et al. 2018; Syme et al. 2018b). Interestingly, the total genome size of the *P*. *teres* f. *teres* isolates ranged from 46.5 Mb to 50.6 Mb (Table 1). Additional assembly metrics are summarized in Table 1 with the currently published *P*. *teres* f. *teres* isolate 0-1 assembly (Wyatt et al. 2018) shown as a reference metric for quality comparison between the four newly sequenced isolates 15A, 6A, FGOH04Ptt-21, and BB25 (Table1).

### Genome annotation

To compare gene content between the five *P. teres* f. *teres* isolates we generated high confidence gene models using RNA sequencing data from each isolate. The total number of annotated genes ranged from 11,551 (6A) to 12,183 (15A) gene models between the five *P*. *teres* f. *teres* isolates with RNAseq evidence for the gene models ranging from 74.2% (0-1) to 98.8% (FGOH04Ptt-21). *P. teres* f. *teres* isolates 0-1, 15A, and 6A were indexed, pooled, and sequenced on a single NextSeq run and isolates FGOH04Ptt-21 and BB25 were indexed, pooled, and sequenced on an additional NextSeq run. Differences in RNAseq coverage between isolates is likely due to the increased depth of sequencing for isolates FGOH04Ptt-21 and BB25. The high number of genes with RNAseq evidence in FGOH04Ptt-21 and BB25 helped to refine gene models in the genome annotations of 0-1, 15A, and 6A as the full set of gene models for all *P. teres* f. *teres* isolates were used to refine gene models for each individual *P. teres* f. *teres* genome. BUSCO analysis runs on each of the *P. teres* f. *teres* genomes using the Ascomycota gene set identified complete genes for 97.4% of the genes for isolate 0-1, 98.1% for isolate 15A, 96.1% for isolate 6A, 98.0% for isolate FGOH04Ptt-21, and 97.8% for isolate BB25. Consistent with genome annotations of other fungal pathogens, many predicted genes do not encode known pfam domains with the number of genes encoding a pfam protein domain ranging between 60.6% (15A) and 62.2% (6A and BB25) (Table2).

Secreted proteins and secondary metabolites are important in fungal biology, specifically with how fungi interact with their environments. Plant pathogens employ these molecules as virulence factors and we therefore identified both the genes encoding secreted proteins and biosynthetic gene clusters. The total number of predicted secreted proteins for the five *P*. *teres* f. *teres* isolates ranged from 1,036 (6A) to 1,066 (15A) (Table 2). Total biosynthetic gene clusters identified in the five *P. teres* f. *teres* isolates ranged from 91 (0-1) to 101 (6A) (Table 2). Relatively consistent numbers of each class of biosynthetic gene clusters were detected in each of the five *P. teres* f. *teres* isolates (Table 2).

**Table 2.**
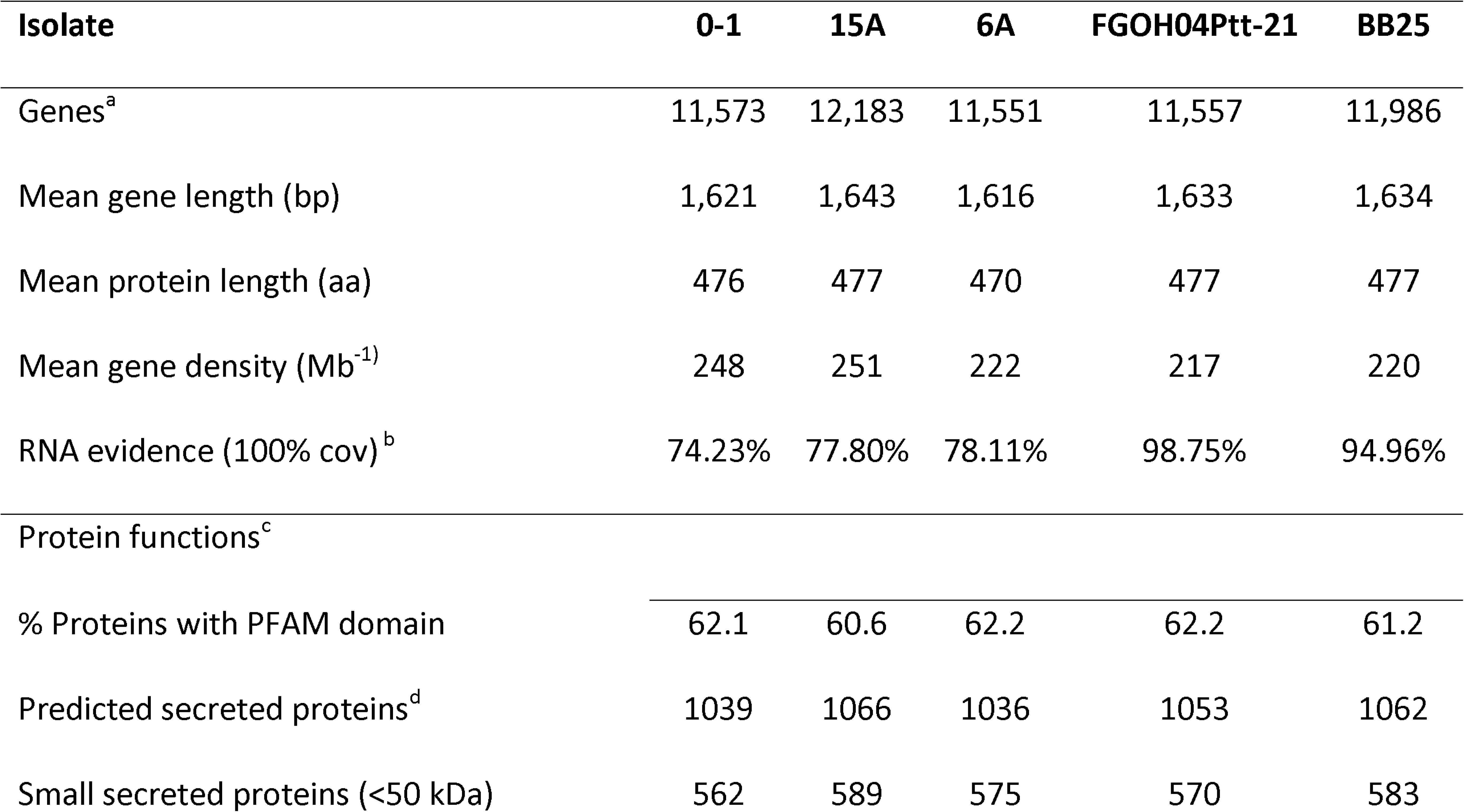
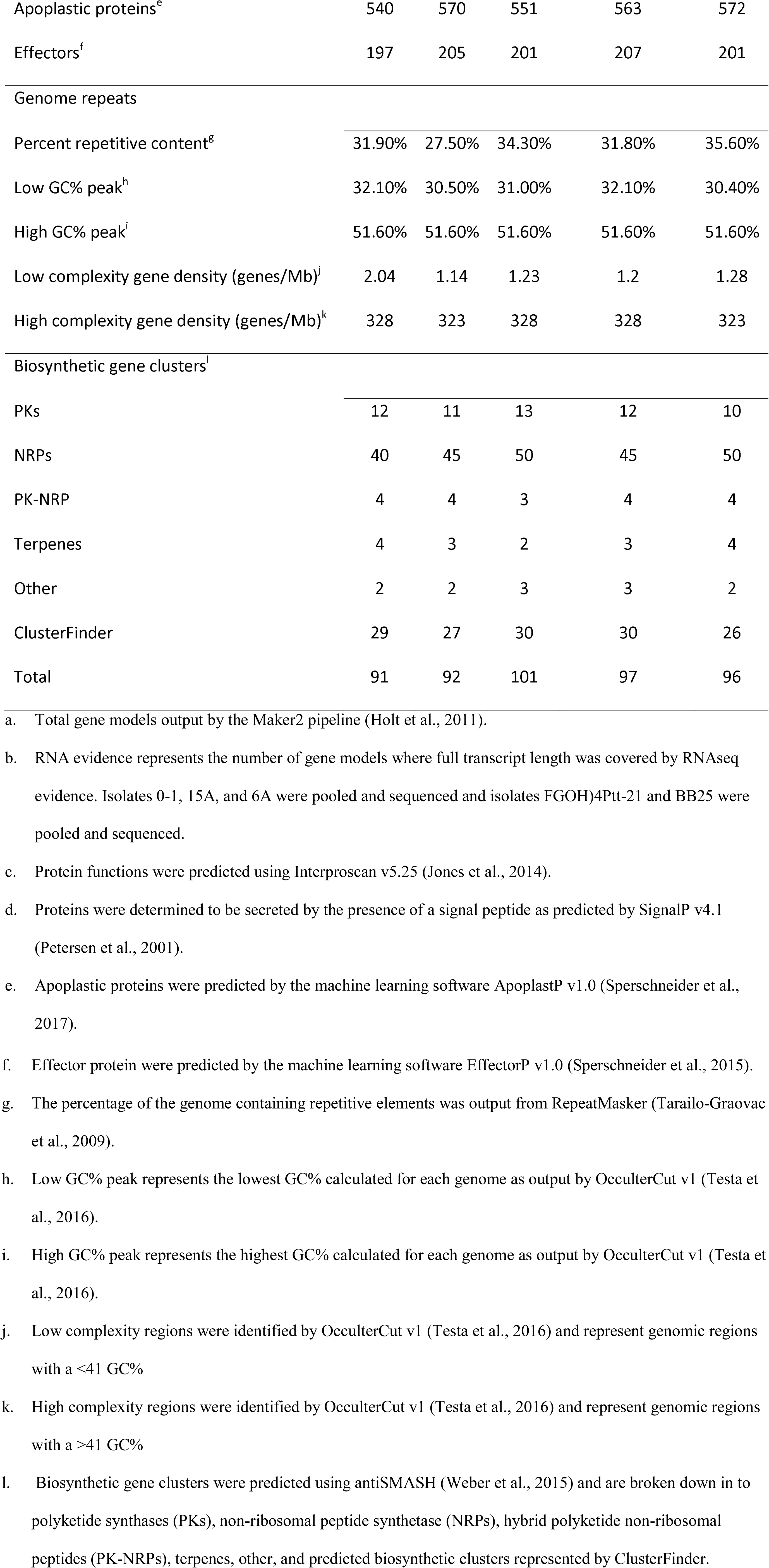
Summary statistics for P. teres f. teres genome annotations.

Annotated repetitive content of the five *P*. *teres* f. *teres* genomes ranged from ∼27.5% (15A) to ∼35.6% (BB25) of the genome (Table 2). Analysis using Occultercut v1 (Testa et al. 2016) revealed a bipartite genome structure divided into high complexity, gene dense regions and low complexity, gene sparse regions (Figure 1). Gene density in low complexity regions ranged from 1.14 genes/Mb to 2.04 genes/Mb and gene density in high complexity regions ranged from 323 genes/Mb to 328 genes/Mb (Table 2). A previous examination of the repeat content of *P. teres* f. *teres* found the most numerous identifiable transposable element families to be the DNA transposable element Tcl-Mariner and the long terminal repeat (LTR) Gypsy element, though a large proportion of annotated repeats belonged to yet unclassified repeat families (Syme et al. 2018b, Wyatt et al. 2018). Repeat elements were also annotated and compared for the five *P. teres* f. *teres* isolates in this study. Overall, the results of the repeat analysis in this study was similar to that of Syme et al. (2018b) in finding the largest proportion of annotated transposable elements belonging to either the DNA Tcl-Mariner transposable element or the LTR-Gypsy element with the LTR-Gypsy element comprising the largest proportion in each of the five *P. teres* f. *teres* genomes (Figure 2). A large proportion of the annotated repeats in the five *P. teres* f. *teres* isolates examined remain unclassified as was also previously noted by Syme et al. (2018b) (Figure 2). Numbers of annotated repetitive elements in the five examined genomes were consistent although isolate 15A had consistently lower numbers of annotated repeats likely due to the lower quality of assembly relative to isolates 0-1, 6A, FGOH04Ptt-21, and BB25 (Figure 2; Table 1).

**Figure 1:**
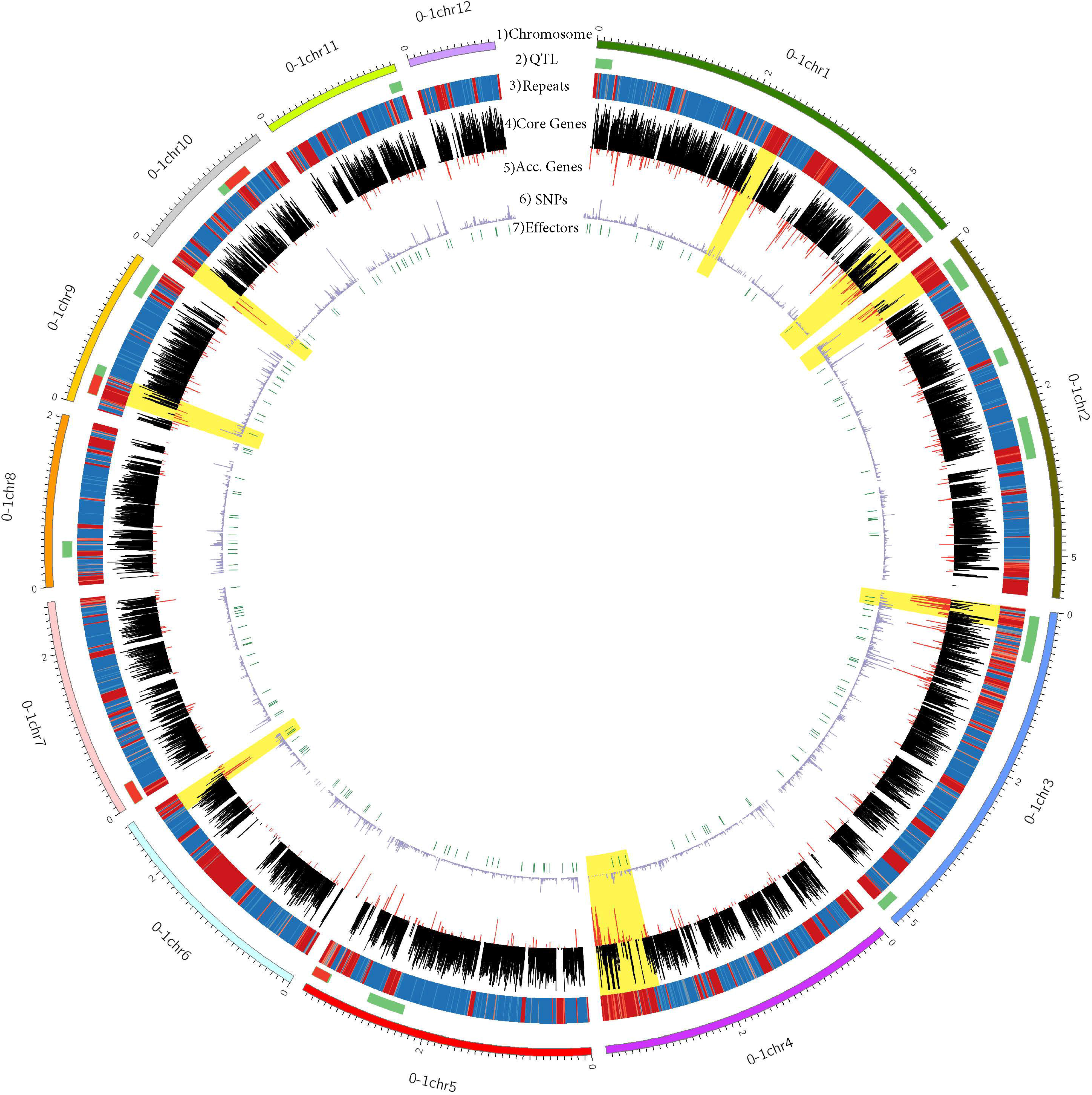
Circos plot showing the genomic landscape of *P. teres* f. *teres* isolate 0-1. **1**) The outermost track of the Circos plot represents the scaffolded chromosomes of isolate 0-1. **2**) The second track shown in green represents published locations of QTL (Lai et al., 2007; Sherve et al., 2014; Koladia et al., 2017). When multiple QTL are detected at a single locus, red is used to mark the overlapping region. **3**) The third track represents a heatmap of repetitive elements in the 0-1 genome where red indicates more repetitiveness and blue denotes repeat sparse and gene dense regions. **4**) The fourth track of the plot represents locations and density of core genes shown in black. **5**) The fifth track represents the locations and density of accessory genes in the 0-1 genome shown in red. **6**) The sixth track shows the density of SNPs identified between the five *P. teres* f. *teres* isolates **7**) The seventh track shows the location of effectors predicted by EffectorP (Sperschneider et al., 2017) shown in green. Yellow blocks highlight exemplary accessory compartments in the 0-1 genome that show more repetitiveness and an increase in accessory gene content.

**Figure 2:**
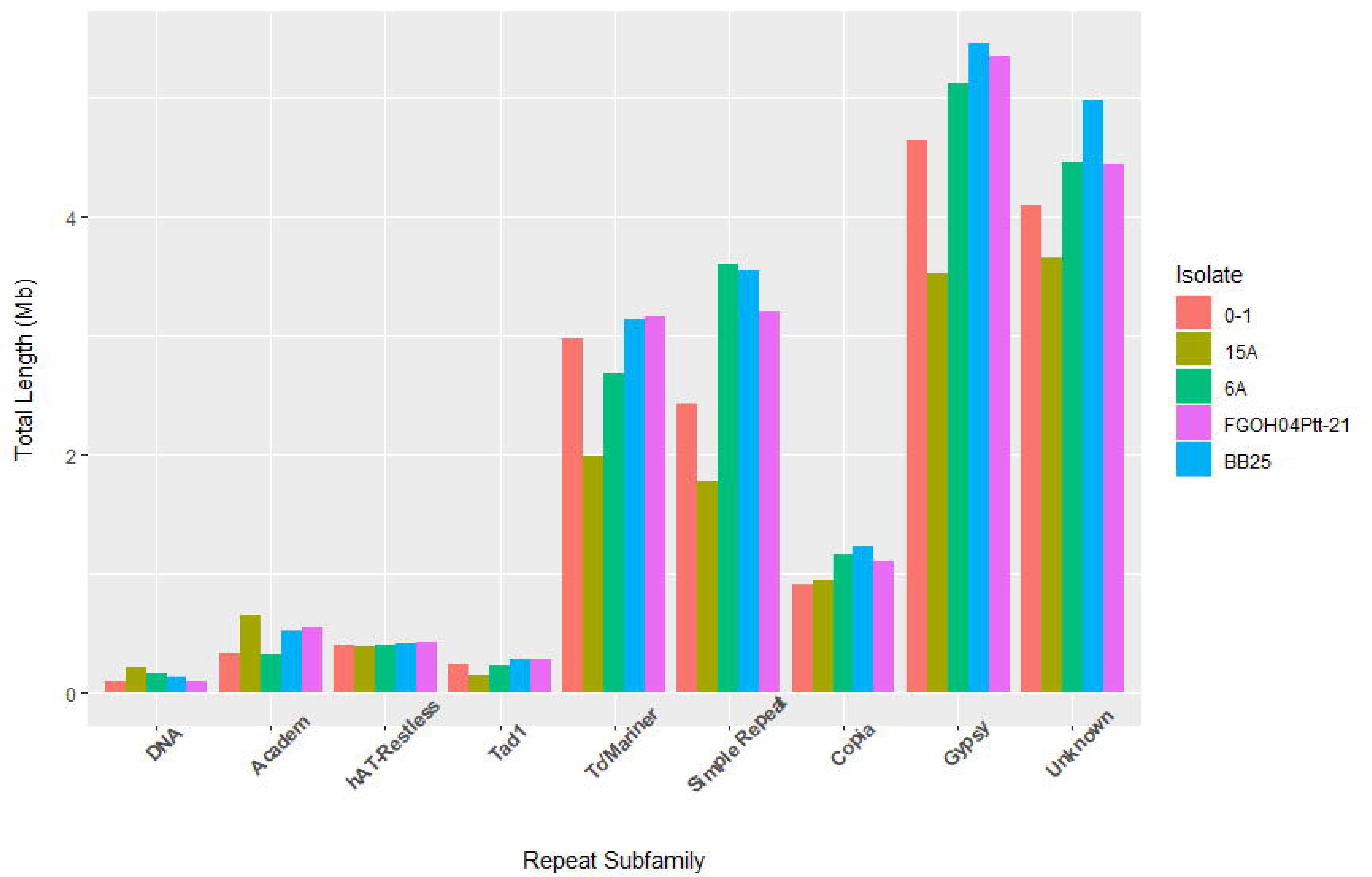
The total length of each type of repeat subfamily expressed in Mbs from each of the five *P. teres* f. *teres* genomes. Amongst all five *P. teres* f. *teres* isolates, the most numerous repeat subfamilies are the Tc/Mariner DNA elements and the Gypsy LTR elements, though a large portion of each genome is comprised of simple repeats and largely unknown/unclassified repetitive elements. *P. teres* f. *teres* isolate 15A shows a reduced number of repeats relative to the other four isolates and this is attributed to uncaptured sequence.

### Comparative pan-genome analysis

To examine macro synteny between *P. teres* f. *teres* isolates, the four newly sequenced isolates 15A, 6A, FGOH04Ptt-21, and BB25 were aligned to the reference 0-1 genome. A high degree of collinearity was observed between the reference isolate 0-1 and isolates 15A, 6A, and FGOH04Ptt-21 along the twelve established chromosomes (Figure 3). The alignment between reference isolate 0-1 and isolate BB25 showed an apparent chromosome fusion between chromosomes 1 and 2 in the BB25 isolate (Figure 4).The fusion of *P. teres* f. *teres* isolate BB25 chromosmome 1 and chromosome 2 also generated a small mini-chromosome comprised of the end of the two chromosomes. It has been previously shown that *P. teres* f. *teres* chromosomes 1 and 2 share synteny along the entire length of *P. tritici-repentis* chromosome 1 (Syme et al. 2018b). Therefore, the genomes of reference isolate 0-1 and BB25 were aligned to the *P*. *tritici-repentis* isolate M4 genome to assess the relatedness of the chromosome 1 and 2 fusion in BB25. The pattern of synteny remained relatively consistent between *P*. *teres* f. *teres* isolates 0-1 and BB25 compared to *P*. *tritici-repentis* isolate M4 (Figure 4). We interpret this to mean that the isolate BB25 chromosome fusion is a recent event and not the result of ancestral inheritance or an interspecies hybridization event. A total of 87,189 SNPs were identified between *P*. *teres* f. *teres* reference isolate 0-1 and the four newly sequenced *P*. *teres* f. *teres* isolates 15A, 6A, FGOH04Ptt-21, and BB25 for use in principal component analysis (Figure 5H). The first principal component accounted for 38% of the variability and the second principal component accounted for 26% of the variability. These results indicate that substantial genetic differentiation exists between isolates, with the exception of 0-1 and FGOH04Ptt-21 clustering closer together.

**Figure 3:**
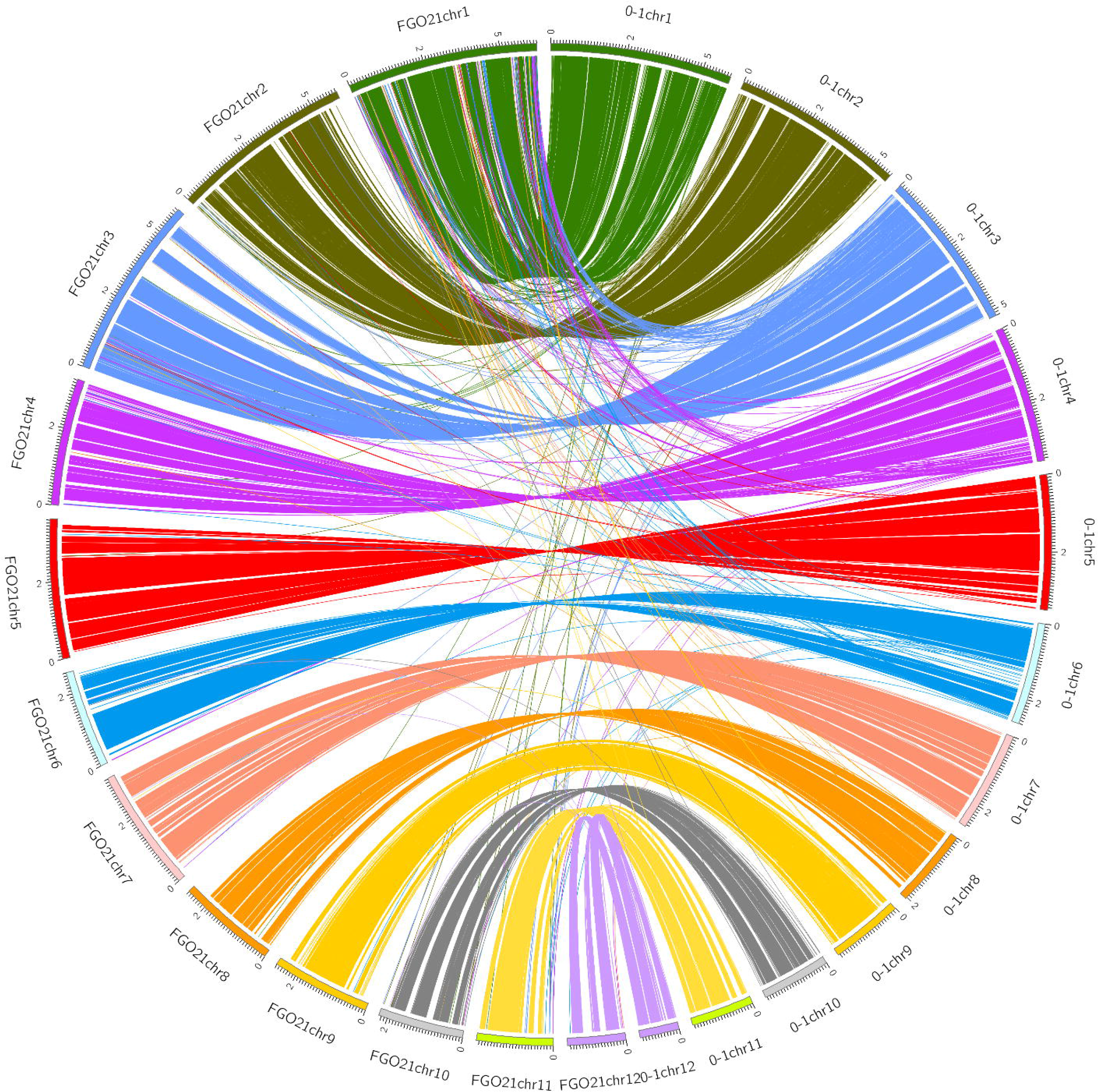
Whole genome synteny plot between *P. teres* f. *teres* isolates 0-1 and FGOH04Pt-21. Bars comprising the outer ring represent individual chromosomes labeled with size (Mb) with tick marks measuring 100 kb. Each ribbon extending from one of the 0-1 chromosomes (0-1chr#) to one of the FGOH04Ptt-21 chromosomes (FGO21chr#) represents a single 0-1 gene’s best hit in the FGOH04Ptt-21 genome. Whole genome synteny plots between *P. teres* f. *teres* isolate 0-1 and isolates 15A, 6A, and BB25 can be found in Supplementary file 3.

**Figure 4:**
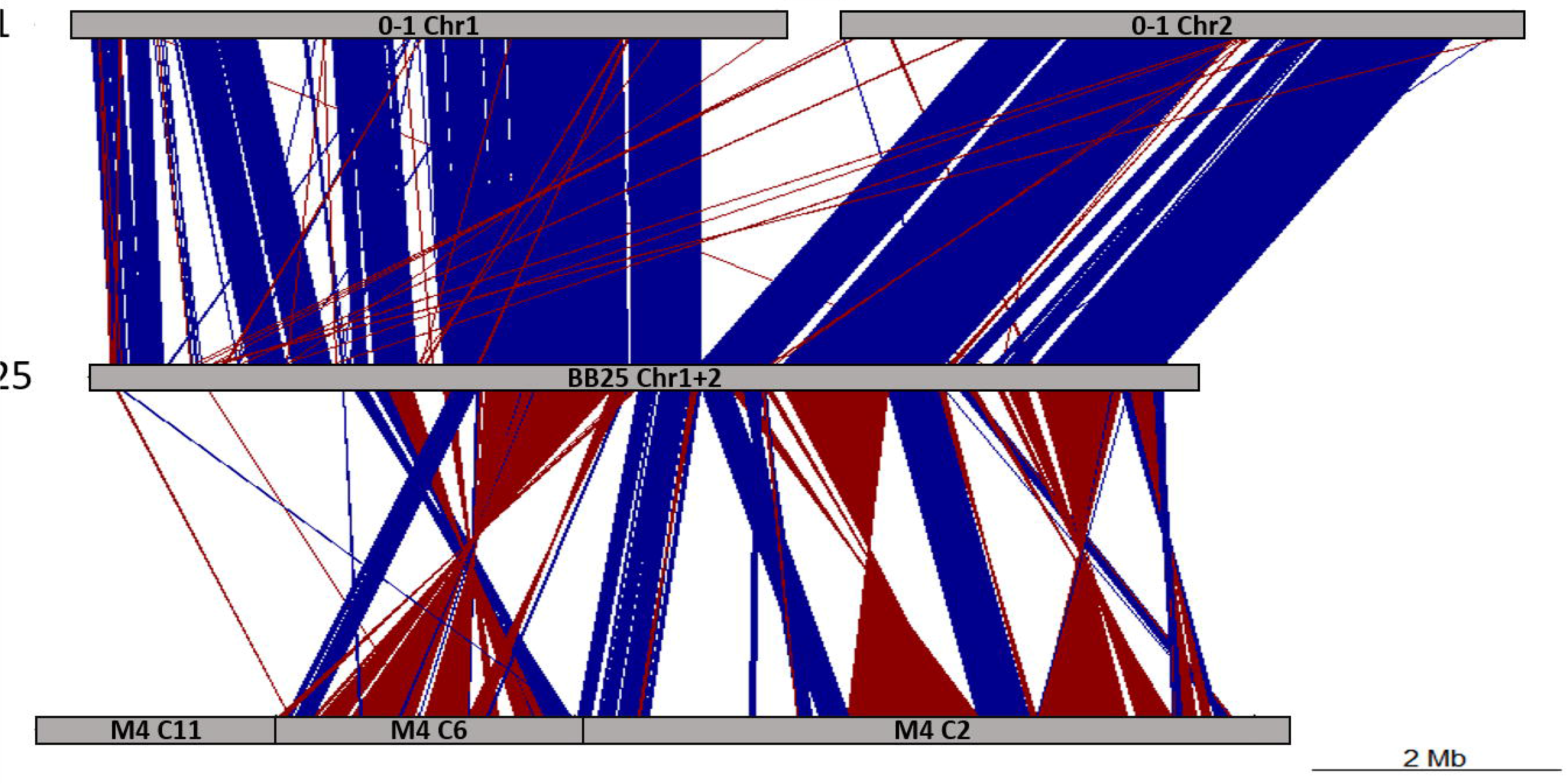
Alignments between *P. teres* f. *teres* isolate 0-1 chromosomes 1 and 2 (top), *P. teres* f. *teres* isolate BB25 fusion chromosome 1-2 (middle), and *P. tritici-repentis* isolate M4 chromosome 1 (bottom). Alignments shown are greater than 2 kb in length and at least 90% identity. Alignments shown in blue represent alignments in the forward orientation and alignments shown in red represent inversions. The pattern of synteny is conserved between the two *P. teres* f. *teres* isolates and both *P. teres* f. *teres* isolates show similar rearrangements relative to *P. tritici-repentis*.

**Figure 5:**
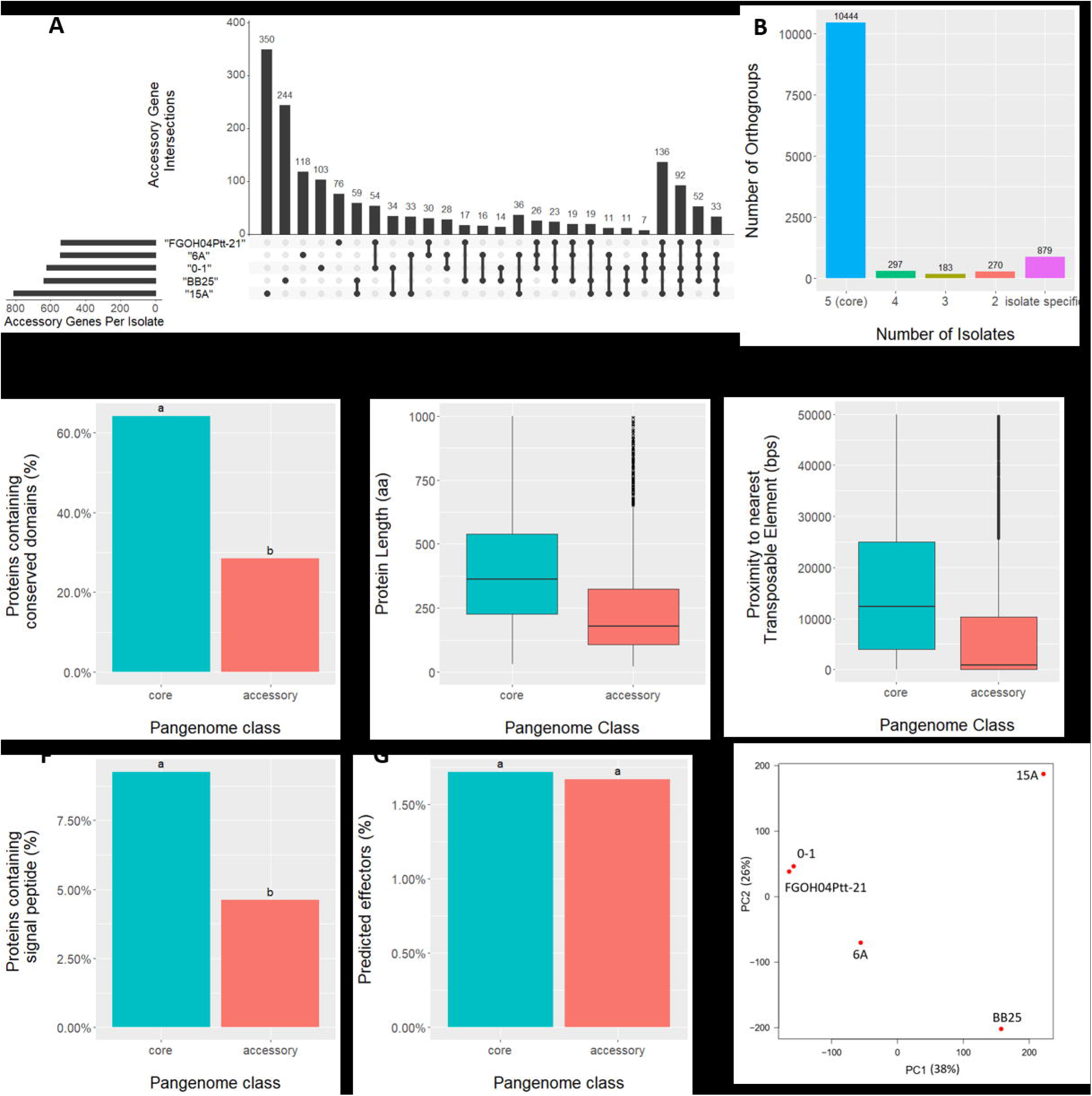
Comparative genomics summary statistics assessed between different gene and protein classes in the *P. teres* f. teres genomes. Panel **A,** is an UpSet diagram representing the number of shared accessory orthogroups between each isolate. The bar graph on top represents the number of proteins in the group and the lower panel indicates which isolates are being compared, with dots indicating the isolate is present and lines drawn between dots to indicate the shared presence. Panel **B,** represents the number of OrthoFinder (Emms *et* al. 2015) determined orthogroups belonging to each isolate. Orthogroups with a protein stemming from each of the five *P. teres* f. *teres* isolates were classified as core proteins and represent the largest group. Orthogroups with proteins stemming from less than five genomes were classified as accessory, and singleton if the orthogroup was derived from just one genome. Panel **C,** shows the different proportions of core and accessory proteins that contain conserved protein domains with letters ‘a’ and ‘b’ above bar plots denoting significantly different groups (Kruskal-Wallis test, P<0.0001). Panel **D,** shows the difference in protein amino acid sequence length between core and accessory proteins (Kruskal-Wallis test, P<0.0001). Panel **E,** shows the difference in the distance to the nearest transposable element (TE) compared between core and accessory proteins (Kruskal-Wallis test, P<0.0001). Panel **F,** shows the different proportions of core and accessory proteins that contain a signal peptide with letters ‘a’ and ‘b’ above bar plots denoting significantly different groups (Kruskal-Wallis test, P<0.0001). Panel **G,** shows the proportion of the core and accessory proteins that are predicted effectors and there was no significant difference found. Panel **H,** shows a principle component analysis that was conducted using >89,156 SNPs.

To assess shared and unique protein functions of the five *P*. *teres* f. *teres* isolates 0-1, 15A, 6A, FGOH04Ptt-21, and BB25, proteins were clustered into homologous families, termed orthogroups, using the program Orthofinder (Emms et al. 2015). The total number of non-redundant protein family orthogroups identified was 12,073. Of the 12,073 orthogroups 10,444 (86.5%) were found to be shared amongst all five *P*. *teres* f. *teres* genomes and constitute the core genome. A total of 10,161 of the 10,444 core orthogroups represented single copy orthogroups with each *P*. *teres* f. *teres* isolate represented, leaving 283 multi-copy orthogroups with shared and unique gene family expansions (Supplementary file 1). Of the 1,629 proteins resulting from the accessory genome, 750 proteins were found to be shared between at least two isolates and 879 proteins were only present in one of the five *P*. *teres* f. *teres* isolates (Figure 5A). The highest number of genes unique to one isolate was observed in 15A (350) and the remaining four *P*. *teres* f. *teres* isolates 0-1, 6A, FGOH04Ptt-21, and BB25 had 103, 118, 76, and 244 unique genes, respectively (Figure 5A).

For further analysis, we subset the pangenome into two categories, the core genome comprised of genes represented in all five genomes and the accessory genome comprised of all genes represented in four or fewer genomes. The average length of amino acid sequence was significantly different between core and accessory genes with core genes encoding longer proteins, 484 amino acids on average, and accessory genes encoding proteins with an average amino acid length of 372 (Kruskal-Wallis test, P<0.0001, Figure 5D). The number of proteins containing a conserved protein domain was also found to be significantly different, with 64.2% of the genes belonging to the core genome encoding proteins containing conserved domains, whereas the proportion of accessory proteins harboring conserved domains was only 28.4% (Figure 5C). Secreted proteins were examined and found to constitute a significantly higher proportion of the core genome when compared to the accessory genome with the core genome containing 9.3% secreted proteins and the accessory genome containing 4.6% secreted proteins (Figure 5F). Predicted effector proteins, as predicted by EffectorP (Sperschider et al. 2016), were also examined and no significant difference in the proportion of encoded effector proteins was found between the core and accessory gene sets with the proportions of each being 1.7% (Figure 5G). Transposable elements have been shown to be important in pathogen genome evolution and were examined for their proximity to core and accessory genes. A significant difference in transposable element proximity was identified with accessory genes clustering nearest transposable elements. On average, transposable elements were found to be 12.6 Kb away from accessory genes and 26.3 Kb away from core genes (Figure 5E).

To assess if core and accessory genomic compartments were evolving at different rates we examined a number of diversity statistics relative to the reference *P. teres* f. *teres* isolate 0-1. We subset the 87,189 SNPs into the relative core and accessory marker sets and found that the mutation rate in accessory genes is slightly elevated at 2.34 SNPs/Kb relative to core genes at 2.04 SNPs/Kb. Additionally, we found that the mutations that do occur in accessory genes are on average twice as likely to be nonsynonymous (missense/silent ratio: 1.3357) relative to the core genome (missense/silent ratio: 0.9168). The calculated pN/pS ratio for core genes was 0.29 and the pN/pS ratio for accessory genes was 0.45 and were significantly different (Kruskal-Wallis test, P<0.0001). Transposable elements were implicated in mutation by insertion into a gene or its promotor, and given the expansion of transposable elements in the *P. teres* f. *teres* genome, we examined the rate of transposable element insertion in genes and their promotor regions between core and accessory gene sets. We found that the proportion of genes in the accessory genome that had a transposable element inserted in the promotor or coding region ranged from24.7% to 36.8% and the core genome ranged from 4.3% to 11.4%. Differences in the type of transposable elements involved in the insertions were observed with LTR-Gypsy, TcMariner-Fot1, hAT-Restless, and DNA-academy transposable elements being inserted in the accessory genome at a higher proportion relative to the core genome.

Gene ontology enrichment analysis was performed on the core and accessory gene sets to further characterize putative functions of the pan-genome. A total of 86 molecular function and 108 biological process gene ontology terms were identified as significantly enriched (p <0.05) within the *P. teres* f. *teres* core genome (Supplementary file 2). Core gene functions showing significant enrichment included basic cellular functions and metabolic processes along with typical housekeeping functions (Supplementary file 2). Gene ontology enrichment analysis of *P*. *teres* f. *teres* accessory genes identified significant enrichments for functions involved in small molecule binding including nucleotide binding, ion binding, and carbohydrate binding. Accessory genes were also enriched in gene ontologies associated with several metabolic processes as well as DNA integration, transposition, and recombination likely driven by the proliferation of transposable elements in the *P. teres* f. *teres* genome detected in the genome annotation (Supplementary file 2).

We identified 283 multi-copy orthogroups representing paralogs of expanded gene families ranging in size from 6 to 203 genes per orthogroup (Table 3; Supplementary file 1). Many of these expanded gene families comprise significantly different gene counts between isolates. For example, the largest family, OG0000000, contains 203 genes with per isolate counts ranging from a single protein in isolate FGOH04Ptt-21 to 104 proteins in isolate 6A (Table 3; Supplementary file 1). Several of the largest orthogroups contain genes with functions in secondary metabolite production and gene functions associated with mechanisms of transposition (Table 3). As an example, the annotated protein domains of OG0000000 included AMP-dependent synthetase/ligase domains, phosphopantetheine binding ACP domains, and condensation domains, often found in proteins involved in the synthesis of polyketides. Annotated protein domains of the second largest and fifth largest groups, OG0000001 and OG0000004, included retrotransposon gag domains, hAT family C-terminal dimerization regions, and ribonuclease H-like domains associated with retro elements. Four orthogroups belonging to a single isolate were identified and represent intragenomic duplication events. The four orthogroups were OG0000214 belonging to isolate 15A and comprising seven genes, OG0000119 belonging to isolate FGOH04Ptt-21 with eleven genes, and orthogroups OG0000098 and OG0000124 belonging to isolate 0-1 and having twelve and ten genes, respectively. Annotated functions of these isolate-specific orthogroups showed functions related to transposition similar to the larger expanded orthogroups.

**Table 3.** Orthogroup summary statistics for the five *P. teres* f. *teres* isolates.

| OrthoGroup | 0-1<br>proteins | 15a<br>proteins | 6a<br>proteins | bb25<br>proteins | FGOH04Ptt-21<br>proteins | Total<br>proteins | Associated functions |
| --- | --- | --- | --- | --- | --- | --- | --- |
| OG0000000 | 3 | 42 | 104 | 53 | 1 | 203 | Secondary metabolite<br>synthesis |
| OG0000001 | 23 | 23 | 17 | 11 | 20 | 94 | Transposable elements |
| OG0000002 | 62 | 11 | 2 | 8 | 4 | 87 | Secondary metabolite<br>synthesis |
| OG0000003 | 31 | 1 | 14 | 10 | 28 | 84 | No known function |
| OG0000004 | 2 | 36 | 24 | 2 | 16 | 80 | Transposable elements |
| OG0000005 | 30 | 5 | 0 | 0 | 41 | 76 | No known function |
| OG0000006 | 0 | 12 | 0 | 2 | 61 | 75 | No known function |
| OG0000007 | 6 | 38 | 2 | 24 | 3 | 73 | No known function |
| OG0000008 | 16 | 15 | 12 | 12 | 11 | 66 | No known function |
| OG0000009 | 5 | 11 | 4 | 33 | 10 | 63 | Transposable elements |
| OG0000010 | 15 | 11 | 10 | 7 | 12 | 55 | No known function |
| OG0000011 | 4 | 20 | 8 | 14 | 9 | 55 | Transposable elements |
| OG0000012 | 11 | 16 | 9 | 10 | 8 | 54 | No known function |
| OG0000013 | 10 | 11 | 9 | 12 | 11 | 53 | No known function |
| OG0000014 | 10 | 16 | 0 | 17 | 10 | 53 | No known function |
| OG0000015 | 10 | 13 | 8 | 11 | 8 | 50 | Epigenetic regulation |

- a. OrthoGroups were output from OrthoFinder (Emms et al., 2015) and groups larger than 50 proteins are shown here. b. Associated functions were interpreted based on results obtained from Interproscan for the genes in each OrthoGroup. OrthoGroups with proteins lacking a conserved domain as predicted by Interproscan were labeled 'No known function'.

### Effector prediction and comparative analysis

Effector proteins are the suite of proteins employed by plant pathogens to manipulate their respective plant hosts and we anticipated the five *P. teres* f. *teres* isolates in this study to have different effector repertoires based on previously published genetic analyses using these isolates (Lai et al. 2007; Shjerve et al. 2014; Koladia et al. 2017). The total number of secreted proteins for each of the five *P*. *teres* f. *teres* isolates ranged from 1,036 secreted proteins (6A) to 1,066 secreted proteins (15A) and, after filtering for a size of less than 50 kDa, the number of small secreted proteins (SSPs) identified ranged from 562 (0-1) to 589 (15A) (Table 2). This set of small secreted proteins was further analyzed using the machine learning programs ApoplastP, designed to predict proteins localized to the apoplast, and EffectorP, designed to predict effector-like proteins. ApoplastP predicted between 95.8-98.7% of SSPs to be localized to the apoplast and EffectorP predicted between 34.4-36.3% of SSPs to have effector-like protein qualities (Table 2). The difference between the proportion of SSPs predicted to be localized to the apoplast versus predicted to be effectors can be attributed to the greater specificity of EffectorP. Not all proteins secreted into the apoplast would be classified as an effector and examples would include the multitude of carbohydrate active enzymes that assist in plant cell wall degradation. Comparative analysis of the set of SSPs for the five *P*. *teres* f. *teres* isolates showed that 2,728 of the 2,879 SSPs belong to the core genome (94.7%) and only 21 SSPs were identified in a single isolate. Of the 21 SSPs unique to a single isolate, eight are predicted to be effectors by EffectorP and eight are predicted to be apoplastic by ApoplastP, however only four of the 21 isolate-specific SSPs are both predicted to be apoplastic and effectors. Protein locus identifiers for the 21 isolate-specific SSPs can be found in the supplementary materials and methods. All 21 isolate-specific SSPs were found to be expressed *in planta* and possibly constitute proteins important for each isolate’s differential response on different barley genotypes.

Comparisons between predicted effector proteins and secreted non-effectors showed significant differences between amino acid sequence length and proximity to transposable elements with predicted effectors having shorter amino acid length and clustering in closer proximity to transposable elements (Kruskal-Wallis P<0.0001). Gene ontology enrichment analysis was performed on the set of SSPs and a total of 56 molecular function and 96 biological process gene ontologies were found to be enriched (p<0.05) (Supplementary file 2). Among the enriched gene ontologies for the set of SSPs were terms associated with hydrolase activity, carbohydrate binding, cell wall degradation and carbohydrate active enzymes (CAZymes) (Supplementary file 2). Gene ontology enrichment analyses was further performed on the set of predicted effector genes to identify any effector specific enrichment. A total of 19 molecular function and 23 biological process gene ontology terms were found to be enriched (Supplementary file 2). Notable among the enriched effector gene ontology terms were hydrolase activity, carbohydrate binding and degradation, regulation of ion transport, and chitin binding. Proteins with hydrolase activity and carbohydrate binding and degradation activity may have important functions in the necrotrophic infection stage of the *P. teres* f. *teres* life cycle and chitin binding proteins have been shown to be involved in protecting and hiding pathogens from plant defense responses (van den Burg et al. 2006; De Jonge et al. 2010; Liu et al. 2016) (Supplementary file 2).

## Discussion

### Genomic Plasticity in P. teres f. teres

In this study we present reference quality genome assemblies of four additional *P*. *teres* f. *teres* isolates including 15A, 6A, FGOH04Ptt-21, and BB25 (Table 1). Each of these isolates have been included in at least one biparental mapping study that investigated the genetics of avirulence/virulence in *P. teres* f. *teres* because of their diverse disease reactions on differential barley lines (Weiland *et* al. 1999; Lai et al. 2007; Shjerve et al. 2014; Koladia et al. 2017). Whole genome comparisons of synteny between the four newly sequenced isolates and the current *P*. *teres* f. *teres* reference sequence 0-1 (Ellwood et al. 2010; Wyatt et al. 2018) revealed evidence of a highly plastic genome structure with conserved synteny among matching chromosomes with large breaks at repeat rich genomic compartments. These repeat rich genomic compartments were common in sub-telomeric regions of the *P. teres* f. *teres* chromosomes (Figure 1, Figure 6). Previous genomic analysis of *P*. *teres* f. *teres* isolates has found more repetitive content relative to the closely related species *P*. *tritici-repentis* and *Parastagonospora nodorum* as well as the other form of the species *P. teres* f. *maculata* (Moolhuijzen et al. 2018; Richards et al. 2018; Syme et al. 2018b). The increase in repetitive content of *P*. *teres* f. *teres* has facilitated a type of genomic fissuring that led to a genomic architecture where regions of highly repetitive DNA are undergoing increased evolutionary speeds relative to the more GC-equilibrated gene-rich regions of the genome (Raffaele and Kamoun 2012; Dong et al. 2015; Faino et al. 2016). Our data, using the five *P. teres* f. *teres* isolates examined in this study, show the importance of repeat dense accessory genomic compartments in the context of previously published disease related quantitative trait loci (QTL). Almost all published *P. teres* f. *teres* disease QTL span accessory compartments and are in sub-telomeric regions of the genome (Figure 1, Figure 6) (Lai et al. 2007; Shjerve et al. 2014; Koladia et al 2017). This makes sense given that these accessory compartments have been shown to be important in the host-pathogen interaction, as they are evolutionary hotspots and often harbor effector proteins (Croll et al. 2012; Thomma et al. 2016; Franceschetti et al. 2017; Seidl et al. 2017; Bertazzoni et al. 2018).

**Figure 6:**
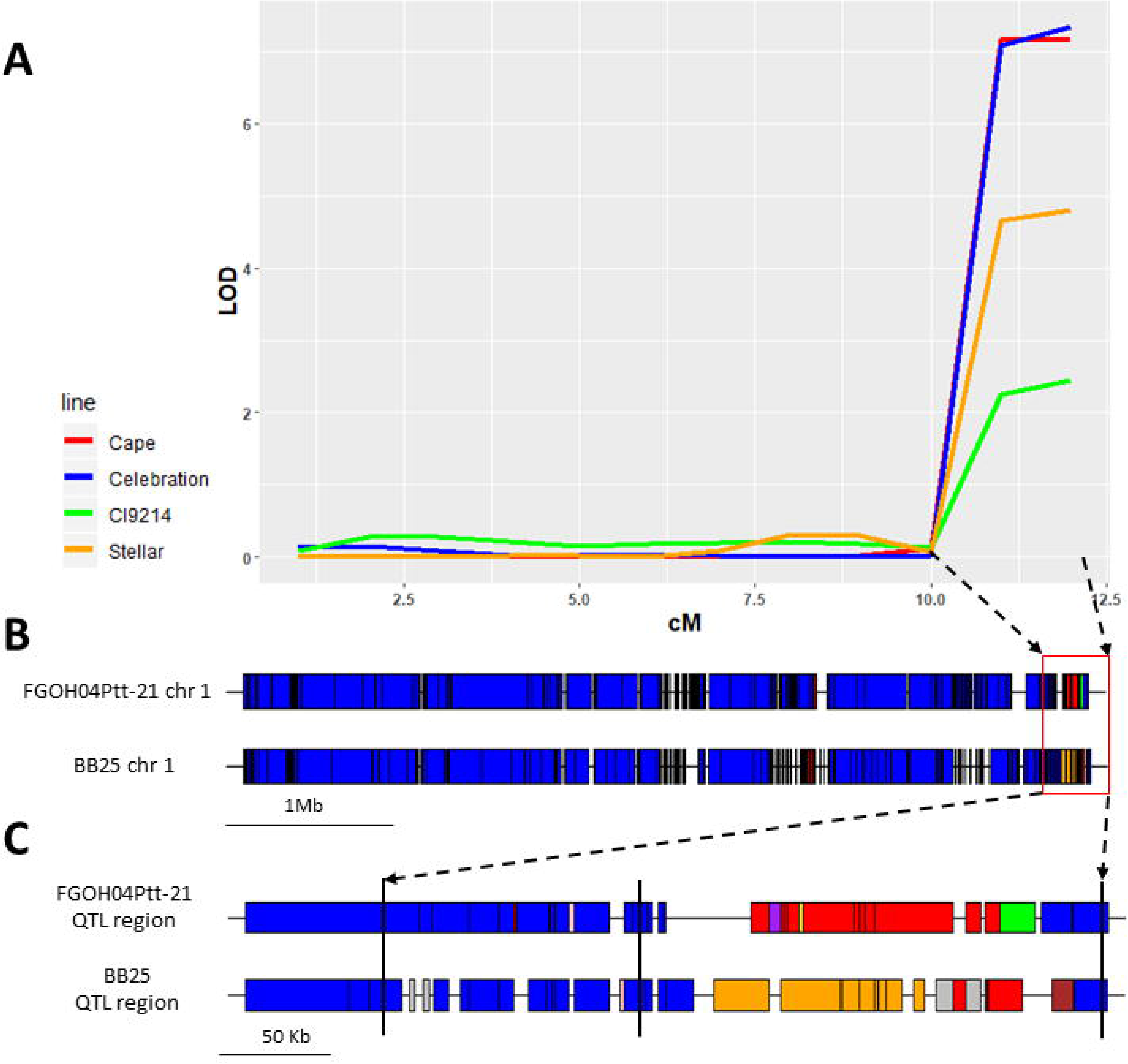
A representative example (Chromosome 1) of nonhomologous chromosomal exchange prevalent at the telomeres of *Pyrenophora teres* f. *teres*. **A**, a quantitative trait locus (QTL) analysis showing a significant association with disease reaction on the barley lines Cape, Celebration, CI9214, and Stellar. LOD score is indicated on the y-axis and centimorgan (cM) distance is indicated on the x-axis. Arrows directed from the QTL region in **A** to the red box in **B** designate the underlying genomic region of *Pyrenophora teres* f. *teres* chromosome 1. **B**, represents the pattern of synteny to the reference *P. teres* f. *teres* isolate 0-1 chromosome 1 along the full length of the chromosome between isolates FGOH04Ptt-21 and BB25. Syntenic regions between matching chromosomes (i.e. chromosome 1 to chromosome 1) are shown in blue. Syntenic regions between non-matching chromosomes are indicated with various colors. Gaps in the alignments are represented by gaps along chromosome 1’s syntenic representation. The red box at the end of the chromosome 1 represents the identified QTL region with black dashed lines with arrows extending to **C** which indicates the genomic region underlying the QTL region identified. **C**, represents the synteny between isolate 0-1 and FGOH04Ptt-21 and BB25, respectively. Vertical black bars indicate the location of the three associated SNP markers that defining the identified QTL region. Synteny between matching chromosomes (i.e. chromosome 1 to chromosome 1) is represented in blue with the different colors representing synteny to sequences not present on isolate 0-1 chromosome 1. Different colors were used to identify different chromosome syntenies. Importantly, the pattern of synteny does not match between isolates FGOH04Ptt-21 and BB25 as is shown by the large red and green block of synteny in isolate FGOH04Ptt-21 and the large orange and red blocks in isolate BB25. Red blocks indicate homology to chromosome 11, green blocks indicate homology to chromosome 2, and orange blocks indicate homology to chromosome 3.

Whole genome alignments were assessed between the five *P*. *teres* f. *teres* isolates to examine the degree of intraspecies synteny and genome plasticity (Figure 3; Supplementary file 3). A chromosome fusion between chromosomes 1 and 2 was discovered in *P*. *teres* f. *teres* isolate BB25 relative to the reference isolate 0-1. Previous work examining whole genome alignments between *P*. *teres* f. *teres* and the closely related species *P*. *tritici-repentis* showed that a similar genome structure was present in *P*. *tritici-repentis* in which *P*. *teres* f. *teres* chromosomes 1 and 2 have synteny along the entire length of *P*. *tritici-repentis* chromosome 1 (Syme et al. 2018b). Isolate BB25’s fused chromosome 1-2 was aligned to the newly published genomic sequence of *P*. *tritici-repentis* isolate M4 (Moolhijzen et al. 2018) to determine if this chromosome fusion was a result of ancestral inheritance. It was found that although *P*. *teres* f. *teres* chromosome 1 and 2 were fused in isolate BB25, the genomic synteny was more conserved between other *P*. *teres* f. *teres* isolates and exhibited the same pattern of rearrangements relative to *P*. *tritici-repentis* (Figure 4). We interpret this to mean that the fusion between chromosome 1 and 2 in *P*. *teres* f. *teres* isolate BB25 to be a recent event and not inherited through ancestry between *P*. *teres* f. *teres* and *P*. *tritici-repentis*. The chromosome fusion in isolate BB25 also created a short chromosome from the remainder of the two fused chromosomes. Unlike mini-chromosomes and accessory chromosomes from other plant pathogen species it does not appear that this short chromosome represents a disposable part of the genome as the sequence content is present in all five *P. teres* f. *teres* isolates. Chromosome length polymorphisms were identified between *P. teres* f. *teres* isolates 0-1 and 15A using pulse field gel electrophoresis (Elwood et al. 2010) and it was recently reported that a chromosome fusion was detected in *P. tritici-repentis* isolate M4 relative to the reference *P. tritici-repentis* isolate Pt-1C-BFP (Moolhuijzen et al. 2018). It is possible that chromosome breakage/fusion events are a common feature of the *Pyrenophora* genus, resulting in chromosome length polymorphisms and chromosome fusion events and represents an interesting area for further investigation.

Chromosome synteny appears relatively conserved along matching chromosomes between *P*. *teres* f. *teres* isolates (Figure 3; Supplementary file 3). Breaks in chromosome collinearity occur primarily around highly repetitive sequences in the *P*. *teres* f. *teres* genomes. Repetitive sequences and genome rearrangements between *P*. *teres* f. *teres* isolates appeared at a higher frequency toward the ends of chromosomes within sub-telomeric regions where genomic synteny was observed to switch between matching chromosomes and all other chromosomes in the genome (Figure 3, Figure 6). A possible mechanism driving these observed structural rearrangements are telomere-centromere breakage-fusion-bridge cycles (Croll et al. 2013) and recombination between non-homologous chromosomes mediated by the high density of repetitive elements that are in accessory genomic compartments, particularly in sub-telomeric regions (Argueso et al. 2008; Bzymek et al. 2001; Raskina et al. 2008).

### Accessory genomic compartments and pathogenicity

Previous genomic work in the closely related species *Zymoseptoria tritici* and *P*. *tritici-repentis* have examined multiple isolates within the same species in a pan-genomic approach to assess the core and accessory gene content of the species (Plissonneau et al. 2018; Moolhuijzen et al. 2018). Core gene content is representative of genes that are present in all members of the species and are likely required for the survival of the species. Accessory gene content includes genes that are absent in members of the same species and are thought to be involved in niche processes. Accessory genomic compartments have been previously shown to undergo rapid evolution relating to high mutation rates, copy number polymorphisms, and frequent ectopic recombination mediated by increased frequency of repetitive elements (Croll et al. 2012). These accessory genomic compartments can be as large as small chromosomes and in the case of *Z. tritici*, include several accessory chromosomes. Though we did not observe accessory chromosomes in *P*. *teres* f. *teres*, there is strong indication of accessory genomic regions within the genome (Figure 1). These accessory genomic regions were frequently identified on the ends of *P. teres* f. *teres* chromosomes where breaks in synteny occur and synteny between non-homologous chromosomes occur. These accessory genomic regions are likely acting in a similar manner to accessory and mini chromosomes found in other fungal species. The *P*. *teres* f. *teres* accessory regions exhibit common characteristics that include high density repetitive content, low density gene content, an increased presence of accessory genes, and the decreased presence of core genes (Figure 1). Accessory genes also exhibited hallmarks of diversifying selection as they had an increased mutational rate and increased pN/pS ratio relative to core genes. Further adding to the increased evolutionary rate of accessory genes was the frequent intersection of accessory genes with transposable elements. The intersection of accessory genes and transposable elements makes sense in *P. teres* f. *teres* given the genomic architecture of accessory genomic compartments described above. In addition to accessory genes clustering in accessory regions, it was observed that a high proportion of secondary metabolite gene clusters were also present in accessory regions of the genome.

Genomic accessory regions and accessory chromosomes have been associated with pathogenicity in multiple plant pathogenic species and genes implicated in virulence have been identified in these regions (Bertazzoni et al. 2018). In the plant pathogen *Leptosphaeria maculans* the *AvrLm11* effector was identified on an accessory chromosome and conferred virulence on *Brassica rapa* cultivars lacking the *Rlm11* gene (Balesdent et al. 2013). The *Magnaporthe oryzae* effector *AvrPik* was also identified on a small 1.6 Mb accessory chromosome (Luo et al. 2007; Chen et al. 2013). To date there have been no published effector genes in *P*. *teres* f. *teres* but there are currently 15 published QTL regions that have been associated with virulence/aviurlence. Of the 15 published QTL, 14 QTL reside in genomic regions that would be categorized as accessory regions and 10 of the 14 QTL are specifically located in subtelomeric regions of the genome (Figure 1, Figure 6) (Weiland et al. 1999; Lai et al. 2007, Shjerve et al. 2014; Koladia et al. 2017). This demonstrates the importance of these rapidly evolving accessory regions in the pathogenicity of *P. teres* f. *teres* (Figure 1, Figure 6). *Gene family expansions in P. teres* f. *teres*

Gene orthology analysis of the five *P. teres* f. *teres* isolates revealed gene family expansions both within single and multiple *P. teres* f. *teres* isolates relative to other *P. teres* f. *teres* isolates examined in this study (Supplementary File 1). Annotated gene functions of expanded gene families within *P. teres* f. *teres* revolved around either, genes related to TEs or, genes relating to the production of secondary metabolites. The association of secondary metabolite synthesis related proteins in the expanded gene families was a common pattern and was observed in other orthologous groups often with a single isolate contributing the majority of the associated homologs (Supplementary file 1). Interestingly, the gene family expansions involving proteins related to secondary metabolite synthesis were present in differing numbers in the five *P. teres* f. *teres* isolates with some clusters being absent from isolates. These secondary metabolite orthogroups may represent niche specific secondary metabolites. Further, QTL regions overlap predicted secondary metabolite clusters pointing to their possible involvement in the disease cycle of *P. teres* f. *teres*. Earlier research on *P. teres* noted the production of phytotoxic molecules that contributed to the disease as they were associated with chlorosis (Smedegard-Petersen 1977; Coval et al. 1990; Bach 1979; Keon and Hargreaves 1983). A recent analysis comparing the number of biosynthetic gene clusters identified between the two forms of *P. teres* (f. *teres* vs. f. *maculata*) using Australian isolates *P. teres* f. *teres* W1-1 and *P. teres* f. *maculata* isolate SG1 found significantly more biosynthetic gene clusters in the *P. teres* f. *teres* isolate (97 total biosynthetic gene clusters) (Syme et al. 2018b). Results of the current study correlate well with that of Syme et al. (2018b) and establish that *P. teres* f. *teres* possesses a wide array of biosynthetic gene clusters (Table 2). *P. teres* f. *teres* biosynthetic gene clusters and their interaction during infection remain an important and unexplored area of research. While further examination of *P. teres* f. *teres* biosynthetic gene clusters is needed, our results confirm that *P. teres* f. *teres* possesses an enormous capacity to produce biologically active secondary metabolites (Syme et al. 2018b; Coval et al. 1990; Smedegard-Petersen 1977).

The second notable observation of the gene orthology analysis was the presence of TE-related proteins in the expanded gene families. Protein domains found in the top expanded gene families included the retrotransposon gag domain, hAT family C-terminal dimerization region, and the ribonuclease H-like domain. The presence of these protein domains in the identified expanded gene families adds support to previous results of annotated repetitive elements in the *P. teres* f. *teres* genome. Previous examinations of the *P. teres* f. *teres* repetitive content showed large repeat expansions relative to *P. teres* f. *maculata* (Syme et al. 2018b).

### Main conclusions and prospects

This study presents four new near complete genome sequences of *P. teres* f. *teres* with gene model annotations and annotated repetitive elements. The four newly sequenced genomes of *P. teres* f. *teres* isolates 15A, 6A, FGOH04Ptt-21, and BB25 were compared to the published reference genome sequence of *P. teres* f. *teres* isolate 0-1 and an intraspecies pan-genomic comparison was performed. Importantly, accessory genomic regions were identified in the *P. teres* f. *teres* genome that represent evolutionarily active genome compartments. Accessory genomic regions represented breaks in genome synteny and held higher numbers of accessory genes and transposable elements. This analysis has set the foundation for understanding the genomic landscape of *P. teres* f. *teres* and the mechanisms involved in this pathogen’s evolution. These resources will also facilitate future research in the barley-*P. teres* f. *teres* pathosystem providing a foundation for future research examining effector biology, secondary metabolites, transposable elements, and evolutionary population genomics.

## Materials and Methods

### Biological materials

Five *P*. *teres* f. *teres* isolates were used in this study, each of which has been included as a parent in a previous virulence mapping study. Isolate 0-1 is the currently published reference genome (Ellwood et al. 2010; Wyatt et al. 2018) and is a Canadian isolate collected in Ontario, Canada (Weiland et al. 1999). Isolates 15A (Steffenson and Webster 1992) and 6A (Wu et al. 2003) were collected in California, USA. Isolate FGOH04Ptt-21 was collected in Fargo, North Dakota, USA (Koladia et al. 2017) and isolate BB25 is an isolate collected in Denmark (kindly provided by Lise Nistrup Jorgensen). These isolates were chosen as a result of their inclusion in previous genetic studies as they differ in virulence and avirulence across barley lines commonly used in differential screening sets. Isolates 15A and 0-1 were first used in a bi-parental mapping study due to their differential reactions on Harbin barley (Weiland *et al*. 1999) and were later used in a bi-parental mapping study examining the differential reactions of these two isolates on the barley lines Tifang, and Prato (Lai *et al*. 2007). Isolate 15A was also crossed to isolate 6A to generate a bi-parental mapping population due to their different disease reactions on Kombar and Rika barley, with isolate 15A being virulent on Kombar but avirulent on Rika and isolate 6A being avirulent on Kombar but virulent on Rika (Sherve *et al*. 2014). Isolates FGOH04Ptt-21 and BB25 were crossed to create a bi-parental mapping population based on the isolates differential disease responses (Koladia *et al*, 2017) on barley lines Manchurian, Tifang, CI4922, Beecher, Celebration, Pinnacle, Hector, and Stellar (Koladia *et al*. 2017).

Fungal tissue for DNA extraction was grown for *P*. *teres* f. *teres* isolates 15A, 6A, FGOH04Ptt-21, and BB25 as described in Koladia et al. 2017 (Supplementary Materials and Methods). High molecular weight DNA extractions were completed following the methods outlined in Richards et al. 2018 (Supplementary Materials and Methods).

### Genome sequencing, assembly, and scaffolding

PacBio 20 kb size-selected sequencing libraries were prepared for all isolates and then sequenced on a PacBio RSII instrument at the Mayo Clinic Molecular Biology Core (Rochester, MN). A total of nine SMRT cells were sequenced for each of the four *P*. *teres* f. *teres* isolates.

Genome assemblies of the four *P*. *teres* f. *teres* isolates 15A, 6A, FGOH04Ptt-21, and BB25 were assembled in a similar manner to 0-1 (Wyatt et al. 2018). The Canu v1.5 assembler (Koren et al. 2017) was used with raw reads input in FASTQ format for correction, trimming, and assembly. The option ‘genomeSize=46.5m’ was used based on the *P*. *teres* f. *teres* reference isolate 0-1 genome size (Wyatt et al. 2018). Genome polishing of each assembly was done with Pilon v1.21 (Walker et al. 2014) using each isolate’s genome and each isolate’s sequencing reads aligned to the genome in BAM file format. Contigs flagged as potential repeats or flagged as potential non-unique contigs by Canu were manually inspected and excluded.

The genetic linkage map of the bi-parental mapping population of FGOH04Ptt-21 × BB25 (Koladia et al. 2017) was used to scaffold the *P*. *teres* f. *teres* isolates 15A, 6A, FGOH04Ptt-21, and BB25 with the program ALLMAPS (Tang et al. 2015). ALLMAPS takes assembled contigs and generates scaffolds based on coordinates from genetic linkage maps, optical maps, or syntenic maps. Contigs were ordered according to marker order on linkage groups with 100 ‘N’ nucleotides inserted to represent gaps. Assembled contigs were quality checked by mapping reads back to the assembly in order to assess collapsed or expanded repeat regions and potentially misassembled regions by manual inspection of read coverage. Contig collinearity with the linkage map was uniform with the exception of isolate BB25 that contained a chromosome fusion that spanned two linkage groups. Reads were mapped back to the BB25 chromosome fusion in order to assess read depth coverage across the fusion point of chromosomes 1 and 2 for confirmation of the chromosome fusion.

Genome assemblies for *P. teres* f. *teres* isolates 15A, 6A, FGOH04Ptt-21, and BB25 were submitted to NCBI Genbank and can be found under accession numbers VBVL00000000 (15A), VFEN00000000 (6A), VBVN00000000 (FGOH04Ptt-21), and VBVM00000000 (BB25). The reference genome for *P. teres* f. *teres* isolate 0-1 was previously submitted to NCBI Genbank and can be found under accession number NPOS00000000.

### RNA sequencing and genome annotation

RNA sequencing was done using *in-planta* time points of 48 h, 72 h, and 96 h post inoculation and a sample from liquid culture. Liquid culture and *in-planta* samples were collected in three replicates. Libraries were sequenced on an Illumina Nextseq at the USDA-ARS Small Grains Genotyping Center (Fargo, ND) to produce 150 bp single-end reads. Illumina FASTQ files were quality checked and trimmed to remove adapters and low quality sequence (Supplementary Materials and Methods). Trimmed reads were aligned and assembled into transcripts following the protocol described in Pertea et al. (2016). For each of the four *P*. *teres* f. *teres* isolates including 15A, 6A, FGOH04Ptt-21, and BB25, RNAseq reads were aligned to each previously assembled genome (Supplementary Materials and Methods).

Genome annotations were compiled using the Maker2 pipeline (Holt et al. 2011) using the same annotation process used in the generation of the 0-1 reference annotation set (Wyatt et al. 2018) (Supplemental Materials and Methods). To evaluate the quality of each *P*. *teres* f. *teres* isolate’s assembled gene models, RNAseq transcript coverage was analyzed with BEDtools using the ‘coverage’ command (Quinlan et al. 2014). Gene models were determined to have RNAseq evidence if a gene model had 100% transcript coverage. Genome annotations were subjected to BUSCO v3 analysis using the Ascomycota data set to assess annotation completeness (Simão et al. 2015) (Supplementary Materials and Methods).

RepeatModeler v1.0.11 (Smit et al. 2015) was used to *de novo* annotate repetitive elements within the genomes. Repetitive elements were then combined from each of the four genomes (15A, 6A, FGOH04Ptt-21, and BB25) and added to the *P*. *teres* f. *teres* specific repeat elements derived in Wyatt et al. (2018). The RepeatModeler *P*. *teres* f. *teres* repeat library was input into RepeatMasker (Tarailo-Graovac et al., 2009) alongside the current release of Repbase (v22.10) (Bao et al. 2015) to soft mask identified repetitive elements and output a final annotation of repetitive elements identified in the genomes. The “buildSummary.pl” RepeatMasker script was applied to gather summary statistics for downstream analysis of repetitive elements.

### Protein domain prediction, detection of biosynthetic gene clusters, and gene ontology

Interproscan v. 5.25 (Jones et al. 2014) was used to predict protein domains and assign gene ontology (GO) terms for each of the *P*. *teres* f. *teres* isolates 0-1, 15A, 6A, FGOH04Ptt-21, and BB25. GO terms (Ashburner et al. 2000) were examined for enrichment using a hyper geometric test with a false discovery rate cut-off set to 0.05 using the R package topGO (Alexa and Rhnenfugrer 2010). SignalP v4.1 was used to identify potential protein secretion signals under default parameters (Petersen et al. 2011) and TMHMM v2.0 was used to predict protein transmembrane domains (Krogh et al. 2001).

Biosynthetic gene clusters were detected in the five *P. teres* f. *teres* genomes using the online analysis tool antiSMASH3.0 (Weber et al. 2015). Two analyses were run with different parameters as in Syme et al. (2018b). A simple analysis was first run on each of the five *P. teres* f. *teres* isolates using unannotated fasta files as input to antiSMASH3.0. Parameters used in the first analysis included the options to run ‘KnownClusterBlast’, ‘SubClusterBlast’, ‘smCoG analysis’, ‘ActiveSiteFinder’, and ‘whole-genome PFAM analysis’. A second run was done and the analysis options ‘ClusterFinder’ and ‘use ClusterFinder algorithm for BGC border prediction’ added. Parameters for the ‘use ClusterFinder algorithm for BGC border prediction’ included setting the minimum cluster size set to five, the minimum number of biosynthesis-related PFAM domains set to five, and a minimum ‘ClusterFinder’ probability of 80%. Extra features added during the second run included ‘Cluster-border prediction based on transcription factor binding sites (CASSIS)’ and ‘ClusterBlast’ (Syme et al. 2018b). The antiSMASH program reports detected number of polyketides (PKs), non-ribosomal peptides (NRPs), PK-NRP hybrids, terpenes, and other (secondary metabolite-like clusters) resulting from the initial simple analysis. An additional report is also generated for ClusterFinder predicted biosynthetic gene clusters.

### Comparative pan-genome analysis

Whole genome alignments of the scaffolded and repeat masked genome assemblies were facilitated by the MUMmer suite using the internal programs ‘nucmer’ for genome alignments (Delcher et al. 2003; Marcais et al. 2018). The four newly sequenced *P*. *teres* f. *teres* isolates 15A, 6A, FGOH04Ptt-21, and BB25 were aligned to the 0-1 reference genome using the ‘nucmer’ program (options: –mum –mincluster 100 –minmatch 50) followed by filtering of the resulting alignment files with ‘delta-filter’ (options: -q -r) to remove any repetitive alignments. Each *P. teres* f. *teres* genome was subjected to analysis using Occultercut v1 to identify clusters of high GC gene dense regions relative to low GC gene sparse regions after identifying and removing mitochondrial sequences (Testa et al. 2016). SNPs were identified by using the Harvest suite (Treangen et al. 2014) program ‘parsnp’ to align *P*. *teres* f. *teres* isolates 15A, 6A, FGOH04Ptt-21, and BB25 to the reference 0-1 assembly (Wyatt et al. 2018). SNPs were input into a principal component analysis (PCA) for the five genomes using the R software package SNPRelate (Zheng et al. 2012). An additional PCA was done using only SNPs from gene coding regions and the resulting eigen values were similar and are supplied in Supplementary Materials and Methods.

The program SNPGenie was used to analyze the 0-1 genome and annotation to determine the number of synonymous and non-synonymous sites (Nelson *et al*. 2015) and the program snpEff was used to calculate the number of mutations classified as synonymous or non-synonymous. The pN/pS ratio was calculated for both the accessory and core gene sets and imported into R for testing significance using a Kruskal-Wallis test. Accessory and core genes were subset and mutations per gene per Kb was calculated by dividing the total mutations in the accessory genes by the length of the corresponding transcripts. Accessory and core genes were checked for intersecting transposable elements using bedtools ‘intersect’ (Quinilan *et al*. 2014).

Proteins were clustered into families using the program OrthoFinder (Emms et al. 2015). The pan-genome was constructed based on OrthoFinder blastp result comparisons between the five *P*. *teres* f. *teres* proteomes (Emms et al. 2015).

### Effector prediction and comparative analysis

Effectors were predicted under the criteria of being secreted, as predicted by SignalP v4.1 (Petersen et al. 2011), lacking a transmembrane domain as predicted by TMHMM v2.0 (Krogh et al. 2001), and having a molecular mass less than 50 kDa. The resulting list of small secreted proteins (SSP) was subjected to further analysis.

Further analyses were run on the set of small secreted proteins, including effector prediction using EffectorP v1.0 (Sperschneider et al. 2015), a program that incorporates machine learning to predict fungal effectors from the secretomes using the default cutoff value. ApoplastP v1.0 (Sperschneider et al. 2017) was also run on the set of SSPs using the default cutoff value. ApoplastP v1.0 is a program that uses machine learning to predict apoplastic localized proteins from non-apoplastic localized proteins. SSPs were further examined for homology both between isolates and within each isolate’s genome. This was accomplished by using ncbi-blast+ ‘blastp’ of the amino acid sequences of each *P*. *teres* f. *teres* SSP set and protein clustering by the program Orthofinder (Emms et al. 2015).

### Data Visualization

Circos plots for Figure 1, Figure 3, and Supplementary Figure 3 were generated on a local Linux server using perl version 5.24. Figure 1 heatmaps were generated with a window size of 5Kb. Figure 1 core and accessory genome frequency plots were graphed as apercent coverage of 5Kb genomic windows. Figure 3 ribbons represent single best matches of genes between *P. teres* f. *teres* isolates.

## Acknowledgments

The authors would like to acknowledge Danielle Holmes for technical assistance. The authors would also like to acknowledge Eva Stukenbrock for valuable suggestions and critical review of the manuscript. This research was supported by the North Dakota Barley Council, National Science Foundation Grant Number: # 1759030, and NIFA-AFRI grant number: #2018-67014-28491.

## Availability of data and materials

*P. teres* f. *teres* genomes and gene models are available at NCBI under BioProject PRJNA434142. Genome assemblies for *P. teres* f. *teres* isolates 15A, 6A, FGOH04Ptt-21, and BB25 were submitted to NCBI Genbank and can be found under accession numbers VBVL00000000 (15A), VFEN00000000 (6A), VBVN00000000 (FGOH04Ptt-21), and VBVM00000000 (BB25). The reference genome for *P. teres* f. *teres* isolate 0-1 was previously submitted to NCBI Genbank and can be found under accession number NPOS00000000. RNA sequencing data was deposited at NCBI SRA and can be found under accession numbers SRR9856883 (0-1 3-day liquid culture), SRR9856882 (0-1 *in-planta* 48h post inoculation), SRR9856885 (0-1 *in-planta* 72h post inoculation), SRR9856884 (0-1 *in-planta* 96h post inoculation); <u>SRR9875069</u> (15A 3 day liquid culture), <u>SRR9875070</u> (15A *in-planta* 48h post inoculation), <u>SRR9875071</u> (15A *in-planta* 72h post inoculation), <u>SRR9875072</u> (15A *in-planta* 96h post inoculation); <u>SRR9875078</u> (6A 3 day liquid culture), <u>SRR9875073</u> (6A *in-planta* 48h post inoculation), <u>SRR9875076</u> (6A *in-planta* 72h post inoculation), <u>SRR9875077</u> (6A *in-planta* 96h post inoculation); <u>SRR9875067</u> (FGOH04Ptt-21 3 day liquid culture), <u>SRR9875068</u> (FGOH04Ptt-21 *in-planta* 48h post inoculation), <u>SRR9875079</u> (FGOH04Ptt-21 *in-planta* 72h post inoculation), <u>SRR9875080</u> (FGOH04Ptt-21 *in-planta* 96h post inoculation); <u>SRR9875074</u> (BB25 3 day liquid culture), <u>SRR9875075</u> (BB25 *in-planta* 48h post inoculation), <u>SRR9875065</u> (BB25 *in-planta* 72h post inoculation), <u>SRR9875066</u> (BB25 *in-planta* 96h post inoculation).

## eXtras

**Supplementary file 1:** OrthoGroups represent groups of orthologous proteins clustered between the five P. teres f. teres genomes produced by Orthofinder (Emms et al. 2015). The left column represents a unique identifier generated by Orthofinder and each column represents the number of proteins from each isolate clustering into the respective orthogroup. The far right column represents the total number of proteins clustering into each orthogroup.

**Supplementary file 2:** Gene ontology enrichment (GOE) results generated for different protein subclasses of the P. teres f. teres genome. Protein subclasses include core proteins, accessory proteins, secreted non-effector proteins, and effector proteins. Each protein catagory was evaluated for GOE for the moleculat function (MF) and biological process (BP) catagories. Outputs were generated with TopGO (Alexa et al. 2019). Gene Ontology ID is reported (GO.ID) with the common term for the Gene Ontology ID reported in the following column. Evalues are reported in the final column and only those greater than 0.05 are reported in this analysis.

**Supplementary file 3:** whole genome synteny plot between P. teres f. teres isolates 0-1 and 15A, 6A and BB25. Bars comprising the outer ring represent individual chromosomes labeled with size (Mb) with tick marks measuring 100 kb. Each ribbon extending from one of the 0-1 chromosomes (0-1chr#) to the corresponding best alignment in the other P. teres f. teres geneomes. Each ribbon represents a >5 Kb syntenic region.

**Supplementary file 4:** Suppary information for the set of 21 small secreted proteins (SSPs) identified as unique to a single isolate of the five P. teres f. teres isolates in this study. The first column lists the geneID, the second column identifies the isolate the SSP was identified in, the third and fourth column show if the gene was identfied by EffectorP or ApoplastP as belonging to those groups respectively (Y=yes, N=No).

**Supplementary materials and methods:** Additional in-depth methods regarding extraction of high molecular weight DNA, tissue collection for RNAseq, RNA sequencing and processing, as well as genome annotation,

## References

Alexa A, Rahnenfuhrer J. 2010. topGO: enrichment analysis for gene ontology. R package version. 2(0).

Andrews S. 2016. FastQC: a quality control tool for high throughput sequence data. Available online at: http://www.bioinformatics.babraham.ac.uk/projects/fastqc

Argueso, J. L., Westmoreland, J., Mieczkowski, P. A., Gawel, M., Petes, T. D., and Resnick, M. A. 2008. Double-strand breaks associated with repetitive DNA can reshape the genome. Proc. Natl. Acad. Sci. U.S.A. 105:11845–11850.

Ashburner, M., Ball, C. A., Blake, J. A., Botstein, D., Butler, H., Cherry, J. M., Davis, A. P., Dolinski, K., Dwight, S. S., Eppig, J. T., and Harris, M. A. 2000. Gene Ontology: tool for the unification of biology. Nat. Genet. 25:25.

Bach, E., Christensen, S., Dalgaard, L., Larsen, P. O., Olsen, C. E., Smedegård-Petersen, V. 1979. Structures, properties and relationship to the aspergillomarasmines of toxins produced by *Pyrenophora teres*. Physiol. Plant Pathol. 14:41–46.

Balesdent, M. H., Fudal, I., Ollivier, B., Bally, P., Grandaubert, J., Eber, F., Chèvre, A. M., Leflon, M., and Rouxel, T. 2013. The dispensable chromosome of *Leptosphaeria maculans* shelters an effector gene conferring avirulence towards *Brassica rapa*. New Phytol. 198:887–898.

Bao, W., Kojima, K. K., and Kohany, O. 2015. Repbase Update, a database of repetitive elements in eukaryotic genomes. Mob. DNA 6:11.

Beattie, A. D., Scoles, G. J., and Rossnagel, B. G. 2007. Identification of molecular markers linked to a *Pyrenophora teres* avirulence gene. Phytopathology 97:842–849.

Bertazzoni, S., Williams, A., Jones, D. A., Syme, R. A., Tan, K. C., and Hane, J. K. 2018. Accessories make the outfit: accessory chromosomes and other dispensable DNA regions in plant-pathogenic Fungi.Mol. Plant Microbe Interact. 31:779–788.

Bialas, A., Zess, E. K., De la Concepcion, J. C., Franceschetti, M., Pennington, H. G., Yoshida, K., Upson, J. L., Chanclud, E., Wu, C.H., Langner, T., and Maqbool, A. 2017. Lessons in effector and NLR biology of plant-microbe systems. bioRxiv171223.

Bolger, A. M., Lohse, M. and Usadel, B. 2014. Trimmomatic: a flexible trimmer for Illumina sequence data. Bioinformatics, 30:2114–2120.

Bzymek, M. and Lovett, S. T. 2001. Instability of repetitive DNA sequences: the role of replication in multiple mechanisms. Proc. Natl. Acad. Sci. U.S.A. 98:8319–8325.

Chen, C., Lian, B., Hu, J., Zhai, H., Wang, X., Venu, R. C., Liu, E., Wang, Z., Chen, M., Wang, B., and Wang, G. L. 2013. Genome comparison of two *Magnaporthe oryzae* field isolates reveals genome variations and potential virulence effectors. BMC genomics, 14:887.

Cingolani, P., Platts, A., Wang, L. L., Coon, M., Nguyen, T., Wang, L., Land, S. J., Lu, X., and Ruden, D. M. 2012. A program for annotating and predicting the effects of single nucleotide polymorphisms, SnpEff: SNPs in the genome of *Drosophila melanogaster* strain w1118; iso-2; iso-3. Fly 6:80–92.

Cook, D. E., Mesarich, C. H., and Thomma, B. P. 2015. Understanding plant immunity as a surveillance system to detect invasion. Annu. Rev. Phytopathol. 53:541–563.

Coval, S. J., Hradil, C. M., Lu, H. S., Clardy, J., Satouri, S., and Strobel, G. A. 1990. Pyrenoline-A and-B, two new phytotoxins from *Pyrenophora teres*. Tetrahedron Lett. 31:2117–2120.

Croll, D. and McDonald, B. A. 2012. The accessory genome as a cradle for adaptive evolution in pathogens. PLoS Pathog. 8:e1002608.

Croll, D., Zala, M., and McDonald, B. A. 2013. Breakage-fusion-bridge cycles and large insertions contribute to the rapid evolution of accessory chromosomes in a fungal pathogen. PLoS Genet. 9:e1003567.

Faino, L., Seidl, M. F., Datema, E., van den Berg, G. C., Janssen, A., Wittenberg, A. H., and Thomma, B. P. 2015. Single-molecule real-time sequencing combined with optical mapping yields completely finished fungal genome. MBio 6:e00936–15.

Delcher, A. L., Salzberg, S. L., and Phillippy, A. M. 2003. Using MUMmer to identify similar regions in large sequence sets. Curr. Protoc. Bioinformatics 1:10–3.

Derbyshire, M., Denton-Giles, M., Hegedus, D., Seifbarghy, S., Rollins, J., van Kan, J., Seidl, M. F., Faino, L., Mbengue, M., Navaud, O., and Raffaele, S. 2017. The complete genome sequence of the phytopathogenic fungus *Sclerotinia sclerotiorum* reveals insights into the genome architecture of broad host range pathogens. Genome Biol. Evol. 9:593–618.

Dong, S., Raffaele, S., and Kamoun, S. 2015. The two-speed genomes of filamentous pathogens: waltz with plants. Curr. Opin. Genet. Dev. 35:57–65.

Ellwood, S. R., Liu, Z., Syme, R. A., Lai, Z., Hane, J. K., Keiper, F., Moffat, C. S., Oliver, R. P., and Friesen, T. L. 2010. A first genome assembly of the barley fungal pathogen *Pyrenophora teres* f. *teres*. Genome Biol. 11:R109.

Emms, D. M. and Kelly, S. 2015. OrthoFinder: solving fundamental biases in whole genome comparisons dramatically improves orthogroup inference accuracy. Genome Biol. 16:157.

Faino, L., Seidl, M. F., Shi-Kunne, X., Pauper, M., van den Berg, G. C., Wittenberg, A. H., and Thomma, B. P. 2016. Transposons passively and actively contribute to evolution of the two-speed genome of a fungal pathogen. Genome Res. 26:1091–1100.

Faris, J. D., Liu, Z., and Xu, S. S. 2013. Genetics of tan spot resistance in wheat. Theor. Appl. Genet. 126:2197–2217.

Franceschetti, M., Maqbool, A., Jiménez-Dalmaroni, M. J., Pennington, H. G., Kamoun, S., and Banfield, M. J. 2017. Effectors of filamentous plant pathogens: commonalities amid diversity. Microbiol. Mol. Biol. Rev. 81:e00066–16.

Friesen, T. L. and Faris, J. D. 2010. Characterization of the wheat-*Stagonospora nodorum* disease system: what is the molecular basis of this quantitative necrotrophic disease interaction? Can. J. Plant Pathol. 32:20–28.

Flor, H.H. 1971. Current status of the gene-for-gene concept. Ann. Rev. Phytopathol. 9:275–296.

Goodwin, S., McPherson, J. D., and McCombie W. R. 2016 Coming of age: ten years of next-generation sequencing technologies. Nat. Rev. Genet. 17:333–351.

Grewal, T. S., Rossnagel, B. G., Pozniak, C. J., and Scoles, G. J. 2008. Mapping quantitative trait loci associated with barley net blotch resistance. Theor. Appl. Genet.116:529–539.

de Guillen, K., Ortiz-Vallejo, D., Gracy, J., Fournier, E., Kroj, T., and Padilla, A. 2015. Structure analysis uncovers a highly diverse but structurally conserved effector family in phytopathogenic fungi. PLoS Patho. 11:e1005228.

Holt, C. and Yandell, M., 2011. MAKER2: an annotation pipeline and genome-database management tool for second-generation genome projects. BMC bioinformatics 12:491.

Hurgobin, B. and Edwards, D. 2017. SNP discovery using a pangenome: has the single reference approach become obsolete? Biology. 6:21.

Jones, J. D. and Dangl, J. L. 2006. The plant immune system. Nature 444:323.

Jones, P., Binns, D., Chang, H. Y., Fraser, M., Li, W., McAnulla, C., McWilliam, H., Maslen, J., Mitchell, A., Nuka, G., and Pesseat, S. 2014. InterProScan 5: genome-scale protein function classification. Bioinformatics 30:1236–1240.

de Jonge, R., van Esse, H. P., Kombrink, A., Shinya, T., Desaki, Y., Bours, R., van der Krol, S., Shibuya, N., Joosten, M. H., and Thomma, B. P. 2010. Conserved fungal LysM effector Ecp6 prevents chitin-triggered immunity in plants. Science 329:953–955.

de Jonge, R., Bolton, M. D., Kombrink, A., van den Berg, G. C., Yadeta, K. A., and Thomma, B. P. 2013. Extensive chromosomal reshuffling drives evolution of virulence in an asexual pathogen. Genome Res. 23:1271–1282.

Van Kan, J. A., Stassen, J. H., Mosbach, A., Van Der Lee, T. A., Faino, L., Farmer, A. D., Papasotiriou, D. G., Zhou, S., Seidl, M. F., Cottam, E., and Edel, D. 2017. A gapless genome sequence of the fungus *Botrytis cinerea*. Mol. Plant Patholo. 18:75–89.

Keon, J. P. R., Hargreaves, J. A. 1983. A cytological study of the net blotch disease of barley caused by *Pyrenophora teres*. Physiol. Plant Pathol. 22:321–IN14.

Khan, T. N. and Boyd, W. J. R. 1969. Environmentally induced variability in the host reaction of barley to net blotch. Aust. J. Biol. Sci. 22:1237–1244.

Kim, D., Langmead, B., and Salzberg, S. L. 2015. HISAT: a fast spliced aligner with low memory requirements. Nat. Methods, 12:357.

Koladia, V. M., Richards, J. K., Wyatt, N. A., Faris, J. D., Brueggeman, R. S., and Friesen, T. L. 2017. Genetic analysis of virulence in the *Pyrenophora teres* f. *teres* population BB25× FGOH04Ptt-21. Fungal Genet. Biol. 107:12–19.

Kombrink, A. and Thomma, B. P. 2013. LysM effectors: secreted proteins supporting fungal life. PLoS Pathog. 9:e1003769.

Koren, S., Walenz, B. P., Berlin, K., Miller, J. R., Bergman, N. H., and Phillippy, A. M. 2017. Canu: scalable and accurate long-read assembly via adaptive k-mer weighting and repeat separation. Genome Res. 27:722–736.

Korf, I. 2004. Gene finding in novel genomes. BMC Bioinformatics 5:59.

Krogh, A., Larsson, B., Von Heijne, G., and Sonnhammer, E. L. 2001. Predicting transmembrane protein topology with a hidden Markov model: application to complete genomes. J. Mol. Biol. 305:567–580.

Krzywinski, M., Schein, J., Birol, I., Connors, J., Gascoyne, R., Horsman, D., Jones, S. J., and Marra, M. A. 2009. Circos: an information aesthetic for comparative genomics. Genome Res. 19:1639–1645.

Lai, Z., Faris, J. D., Weiland, J. J., Steffenson, B. J., and Friesen, T. L. 2007. Genetic mapping of *Pyrenophora teres* f. *teres* genes conferring avirulence on barley. Fungal Genet. Biol. 44:323–329.

Liu, Z., Ellwood, S.R., Oliver, R.P., and Friesen, T.L. 2011. *Pyrenophora teres*: profile of an increasingly damaging barley pathogen. Mol. Plant Pathol. 12:1–19.

Liu, Z., Zhang, Z., Faris, J. D., Oliver, R. P., Syme, R., McDonald, M. C., McDonald, B. A., Solomon, P. S., Lu, S., Shelver, W. L., and Xu, S. 2012a. The cysteine rich necrotrophic effector SnTox1 produced by *Stagonospora nodorum* triggers susceptibility of wheat lines harboring Snn1. PLoS Pathog. 8:e1002467.

Liu, Z. H., Zhong, S., Stasko, A. K., Edwards, M. C., and Friesen, T. L. 2012b. Virulence profile and genetic structure of a North Dakota population of *Pyrenophora teres* f. *teres*, the causal agent of net form net blotch of barley. Phytopathology. 102:539–546.

Liu, Z., Holmes, D. J., Faris, J. D., Chao, S., Brueggeman, R. S., Edwards, M. C., and Friesen, T. L. 2015. Necrotrophic effector triggered susceptibility (NETS) underlies the barley– *Pyrenophora teres* f. *teres* interaction specific to chromosome 6H. Mol. Plant Pathol. 16:188–200.

Liu, Z., Gao, Y., Kim, Y. M., Faris, J. D., Shelver, W. L., Wit, P. J., Xu, S. S., and Friesen, T. L., 2016. SnTox1, a *Parastagonospora nodorum* necrotrophic effector, is a dual function protein that facilitates infection while protecting from wheat produced chitinases. New Phytol. 211:1052–1064.

Luo, C. X., Yin, L. F., Ohtaka, K., and Kusaba, M. 2007. The 1.6 Mb chromosome carrying the avirulence gene AvrPik in *Magnaporthe oryzae* isolate 84R-62B is a chimera containing chromosome 1 sequences. Mycol. Res. 111:232–239.

Manning, V. A., Pandelova, I., Dhillon, B., Wilhelm, L. J., Goodwin, S. B., Berlin, A. M., Figueroa, M., Freitag, M., Hane, J. K., Henrissat, B., and Holman, W. H. 2013. Comparative genomics of a plant-pathogenic fungus, *Pyrenophora tritici-repentis*, reveals transduplication and the impact of repeat elements on pathogenicity and population divergence. G3. 3:41–63.

Marçais, G., Delcher, A. L., Phillippy, A. M., Coston, R., Salzberg, S. L., and Zimin, A. 2018. MUMmer4: A fast and versatile genome alignment system. PLoS Comput. Biol. 14:e1005944.

Mathre, D. E., Kushnak, G. D., Martin, J. M., Grey W. E., and Johnston, R. H. 1997. Effect of residue management on barley production in the presence of Net Blotch Disease. J. Prod. Agric. 10:323–326.

Möller, M. and Stukenbrock, E. H. 2017. Evolution and genome architecture in fungal plant pathogens. Nat. Rev. Microbiol. 15:756.

Moolhuijzen, P., See, P. T., Hane, J. K., Shi, G., Liu, Z., Oliver, R. P., and Moffat, C. S. 2018. Comparative genomics of the wheat fungal pathogen *Pyrenophora tritici-repentis* reveals chromosomal variations and genome plasticity. BMC Genomics. 19:279.

Nelson, C. W., Moncla, L. H. and Hughes, A. L. 2015. SNPGenie: estimating evolutionary parameters to detect natural selection using pooled next-generation sequencing data. Bioinformatics, 31(22), pp.3709–3711.

Perfect, S. E. and Green, J. R. 2001. Infection structures of biotrophic and hemibiotrophic fungal plant pathogens. Mol. Plant Pathol. 2:101–108.

Pertea, M., Pertea, G.M., Antonescu, C. M., Chang, T. C., Mendell, J. T., and Salzberg, S. L. 2015. StringTie enables improved reconstruction of a transcriptome from RNA-seq reads. Nature Biotechnol. 33:290.

Pertea, M., Kim, D., Pertea, G. M., Leek, J. T., and Salzberg, S. L. 2016. Transcript-level expression analysis of RNA-seq experiments with HISAT, StringTie and Ballgown. Nature Protoc. 11:1650.

Petersen, T. N., Brunak, S., von Heijne, G. and Nielsen, H. 2011. SignalP 4.0: discriminating signal peptides from transmembrane regions. Nature Methods. 8:785.

Plissonneau, C., Hartmann, F. E., and Croll, D. 2018. Pangenome analyses of the wheat pathogen *Zymoseptoria tritici* reveal the structural basis of a highly plastic eukaryotic genome. BMC Biol. 16:5.

Quinlan, A. R. 2014. BEDTools: the Swiss army tool for genome feature analysis. Curr. Protoc. Bioinformatics. 11–12.

Raffaele, S., Farrer, R. A., Cano, L. M., Studholme, D. J., MacLean, D., Thines, M., Jiang, R. H., Zody, M. C., Kunjeti, S. G., Donofrio, N. M., and Meyers, B. C. 2010. Genome evolution following host jumps in the Irish potato famine pathogen lineage. Science. 330:1540–1543.

Raffaele, S. and Kamoun, S., 2012. Genome evolution in filamentous plant pathogens: why bigger can be better. Nat. Rev. Microbiol., 10:417.

Raskina, O., Barber, J. C., Nevo, E., and Belyayev, A. 2008. Repetitive DNA and chromosomal rearrangements: speciation-related events in plant genomes. Cytogenet Genome Res. 120:351–357.

Richards, J. K., Wyatt, N. A., Liu, Z., Faris, J. D., and Friesen, T. L. 2018. Reference quality genome assemblies of three *Parastagonospora nodorum* isolates differing in virulence on wheat. G3. 8:393–399.

Rouxel, T., Grandaubert, J., Hane, J. K., Hoede, C., Van de Wouw, A. P., Couloux, A., Dominguez, V., Anthouard, V., Bally, P., Bourras, S., and Cozijnsen, A. J. 2011. Effector diversification within compartments of the *Leptosphaeria maculans* genome affected by Repeat-Induced Point mutations. Nat. Commun. 2:202.

Sánchez-Vallet, A., Fouché, S., Fudal, I., Hartmann, F.E., Soyer, J. L., Tellier, A., and Croll, D. 2018. The Genome Biology of Effector Gene Evolution in Filamentous Plant Pathogens. Ann. Rev. Phytopathol. 56:21–40.

Seidl, M. F. and Thomma, B. P. 2017. Transposable elements direct the coevolution between plants and microbes. Trends Genet. 33:842–851.

Shjerve, R. A., Faris, J. D., Brueggeman, R. S., Yan, C., Zhu, Y., Koladia, V., and Friesen, T. L. 2014. Evaluation of a *Pyrenophora teres* f. *teres* mapping population reveals multiple independent interactions with a region of barley chromosome 6H. Fungal Genet Biol. 70:104–112.

Simão, F. A., Waterhouse, R. M., Ioannidis, P., Kriventseva, E. V., and Zdobnov, E. M. 2015. BUSCO: assessing genome assembly and annotation completeness with single-copy orthologs. Bioinformatics, 31:3210–3212.

Smedegård-Petersen, V. 1977. Isolation of two toxins produced by *Pyrenophora teres* and their significance in disease development of net-spot blotch of barley. Physiol. Plant Pathol. 10:203–211.

Smit, A. F. A. and Hubley, R., 2008. RepeatModeler Open-1.0. Available from http://www.repeatmasker.org.

Sperschneider, J, Gardiner, D. M., Dodds, P. N., Tini, F., Covarelli, L., Singh KB, Manners, J. M., Taylor, J. M. 2015 EffectorP: Predicting Fungal Effector Proteins from Secretomes Using Machine Learning. New Phytol. 210:743–761.

Sperschneider, J., Dodds, P. N., Singh, K. B., and Taylor, J. M. 2017. ApoplastP: prediction of effectors and plant proteins in the apoplast using machine learning. bioRxiv. 182428.

Stahl, E. A. and Bishop, J. G. 2000. Plant–pathogen arms races at the molecular level. Curr. Opin.Plant Biol. 3:299–304.

Stanke, M., Schöffmann, O., Morgenstern, B., and Waack, S. 2006. Gene prediction in eukaryotes with a generalized hidden Markov model that uses hints from external sources. BMC Bioinformatics. 7:62.

Steffenson, B. J. and Webster, R. K. 1992. Pathotype diversity of *Pyrenophora teres* f. *teres* on barley. Phytopathology. 82:170–177.

Syme, R. A., Tan, K. C., Rybak, K., Friesen, T. L., McDonald, B. A., Oliver, R. P., and Hane, J. K. 2018a. Pan-*Parastagonospora* Comparative Genome Analysis—Effector Prediction and Genome Evolution. Genome Biol. Evol. 10:2443–2457.

Syme, R., Martin, A., Wyatt, N. A., Lawrence, J., Muria-Gonzalez, M., Friesen, T. L., and Ellwood, S. 2018b. Transposable element genomic fissuring in *Pyrenophora teres* is associated with genome expansion and dynamics of host-pathogen genetic interactions. Front. Genet. 9:130.

Tang, H., Zhang, X., Miao, C., Zhang, J., Ming, R., Schnable, J. C., Schnable, P. S., Lyons, E., and Lu, J. 2015. ALLMAPS: robust scaffold ordering based on multiple maps. Genome Biol. 16:3.

Tarailo Graovac, M. and Chen, N. 2009. Using RepeatMasker to identify repetitive elements in genomic sequences. Curr. Protoc. Bioinformatics. 5:4–10.

Ter-Hovhannisyan, V., Lomsadze, A., Chernoff, Y. O., and Borodovsky, M. 2008. Gene prediction in novel fungal genomes using an *ab initio* algorithm with unsupervised training. Genome Res. 18:1979–1990.

Testa, A. C., Oliver, R. P., and Hane, J. K. 2016. OcculterCut: a comprehensive survey of AT-rich regions in fungal genomes. Genome Biol. Evol. 8:2044–2064.

Tettelin, H., Masignani, V., Cieslewicz, M. J., Donati, C., Medini, D., Ward, N. L., Angiuoli, S. V., Crabtree, J., Jones, A. L., Durkin, A. S., and DeBoy, R. T. 2005. Genome analysis of multiple pathogenic isolates of *Streptococcus agalactiae*: implications for the microbial “pan-genome”. Proc. Nat. Acad. Sci. 102:13950–13955.

Thomma, B. P., Seidl, M. F., Shi-Kunne, X., Cook, D. E., Bolton, M. D., van Kan, J. A., and Faino, L. 2016. Mind the gap; seven reasons to close fragmented genome assemblies. Fungal Genet Biol. 90:24–30.

Treangen, T. J., Ondov, B. D., Koren, S., and Phillippy, A. M. 2014. The Harvest suite for rapid core-genome alignment and visualization of thousands of intraspecific microbial genomes. Genome Biol. 15:524.

van den Burg, H. A., Harrison, S. J., Joosten, M. H., Vervoort, J., and de Wit, P. J. 2006. *Cladosporium fulvum* Avr4 protects fungal cell walls against hydrolysis by plant chitinases accumulating during infection. Mol. Plant Microbe Interact. 19:1420–1430.

Vernikos, G., Medini, D., Riley, D. R. and Tettelin, H. 2015. Ten years of pan-genome analyses. Curr. Opin Microbiol. 23:148–154.

Vleeshouwers, V. G. and Oliver, R. P. 2014. Effectors as tools in disease resistance breeding against biotrophic, hemibiotrophic, and necrotrophic plant pathogens. Mol. Plant Microbe Interact. 27:196–206.

Walker, B. J., Abeel, T., Shea, T., Priest, M., Abouelliel, A., Sakthikumar, S., Cuomo, C. A., Zeng, Q., Wortman, J., Young, S. K., and Earl, A. M., 2014. Pilon: an integrated tool for comprehensive microbial variant detection and genome assembly improvement. PloS One. 9:e112963.

Weber, T., Blin, K., Duddela, S., Krug, D., Kim, H. U., Bruccoleri, R., Lee, S. Y., Fischbach, M. A., Müller, R., Wohlleben, W., and Breitling, R. 2015. antiSMASH 3.0 - a comprehensive resource for the genome mining of biosynthetic gene clusters. Nucleic Acids Res. 43:W237–W243.

Weiland, J. J., Steffenson, B. J., Cartwright, R. D., and Webster, R. K. 1999. Identification of molecular genetic markers in *Pyrenophora teres* f. *teres* associated with low virulence on ‘Harbin’barley. Phytopathology. 89:176–181.

Wu, H. L., Steffenson, B. J., Zhong, S., Li, Y. and Oleson, A. E. 2003. Genetic variation for virulence and RFLP markers in *Pyrenophora teres*. Can. J. Plant Pathol. 25:82–90.

Wyatt, N. A., Richards, J. K., Brueggeman, R. S., and Friesen, T. L. 2018. Reference assembly and annotation of the *Pyrenophora teres* f. *teres* isolate 0-1. G3. 8:1–8.

Yoshida, K., Saitoh, H., Fujisawa, S., Kanzaki, H., Matsumura, H., Yoshida, K., Tosa, Y., Chuma, I., Takano, Y., Win, J., and Kamoun, S. 2009. Association genetics reveals three novel avirulence genes from the rice blast fungal pathogen *Magnaporthe oryzae*. Plant Cell. 21:1573–1591.

Zheng, X., Levine D., Shen, J., Gogarten, S., Laurie, C., Weir, B. 2012. A high-performance computing toolset for relatedness and principal component analysis of SNP data. Bioinformatics. 28:3326–3328.

Zipfel, C. 2009. Early molecular events in PAMP-triggered immunity. Curr. Opin. Plant Biol. 12:414–420.

