## Supplementary figures and images for "A comparative genomic analysis of the barley pathogen *Pyrenophora teres* f. *teres* identifies sub-telomeric regions as drivers of virulence"

### Supplementary file 3

#### Slide 1
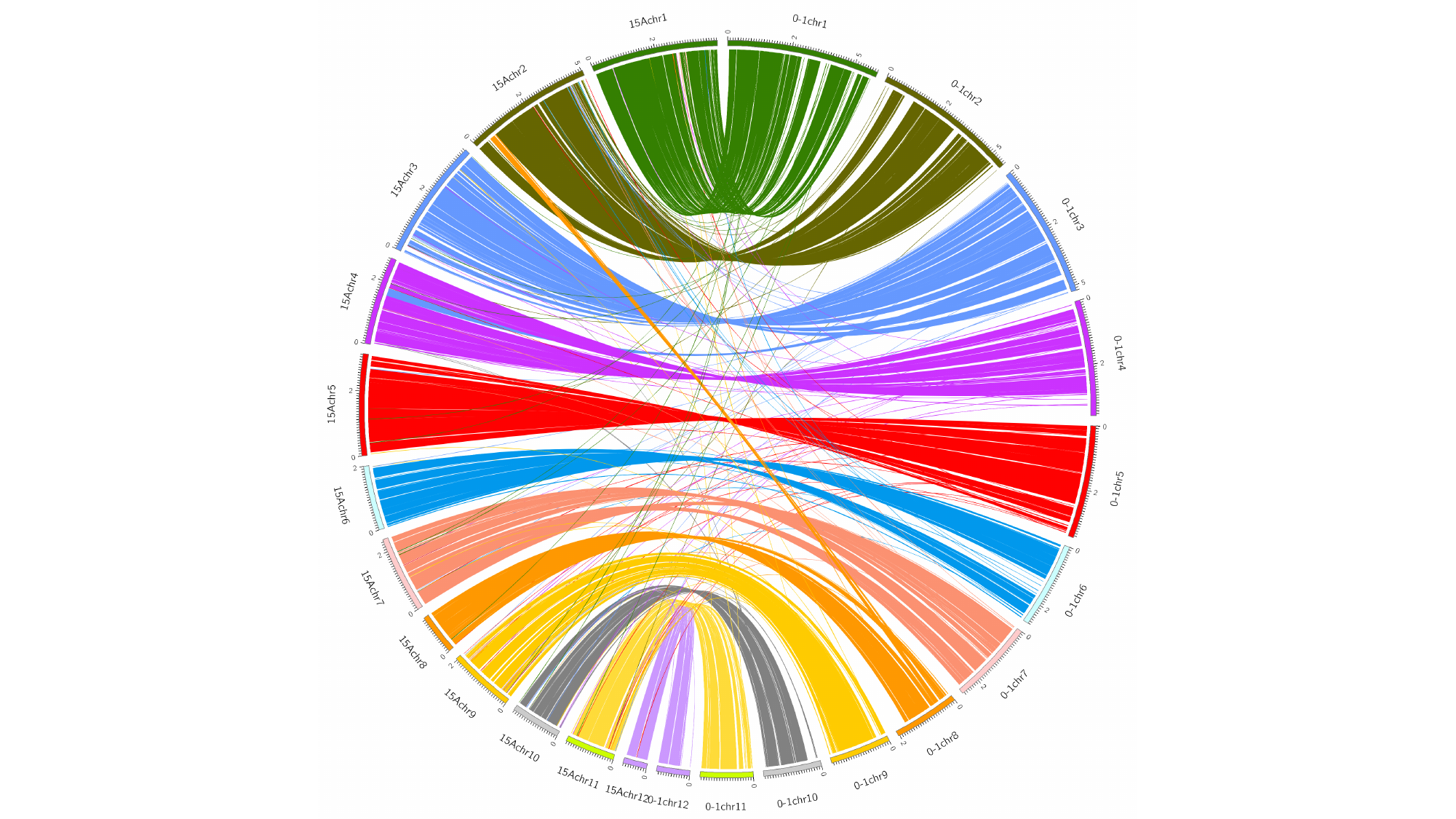

#### Slide 2
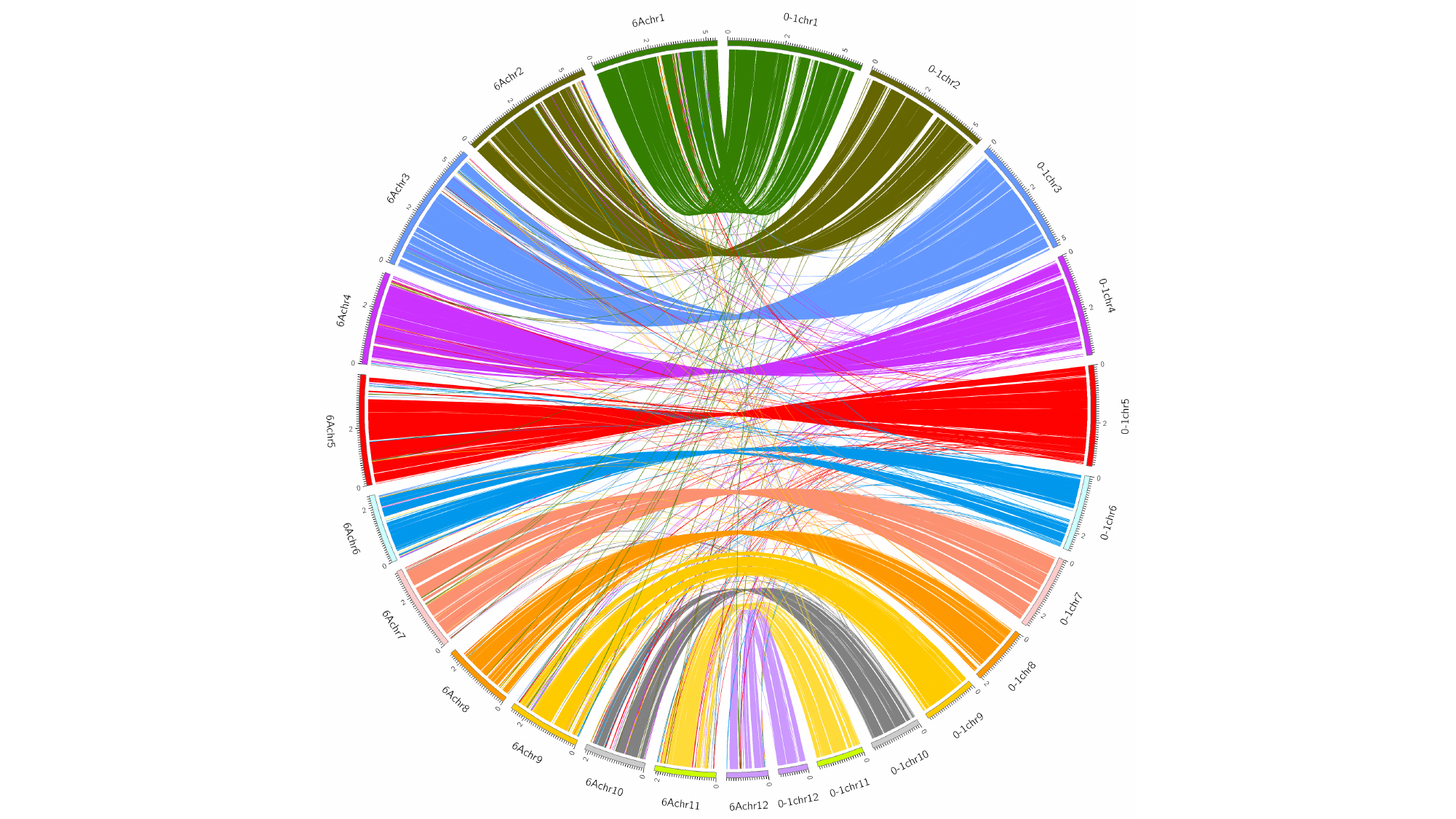

#### Slide 3
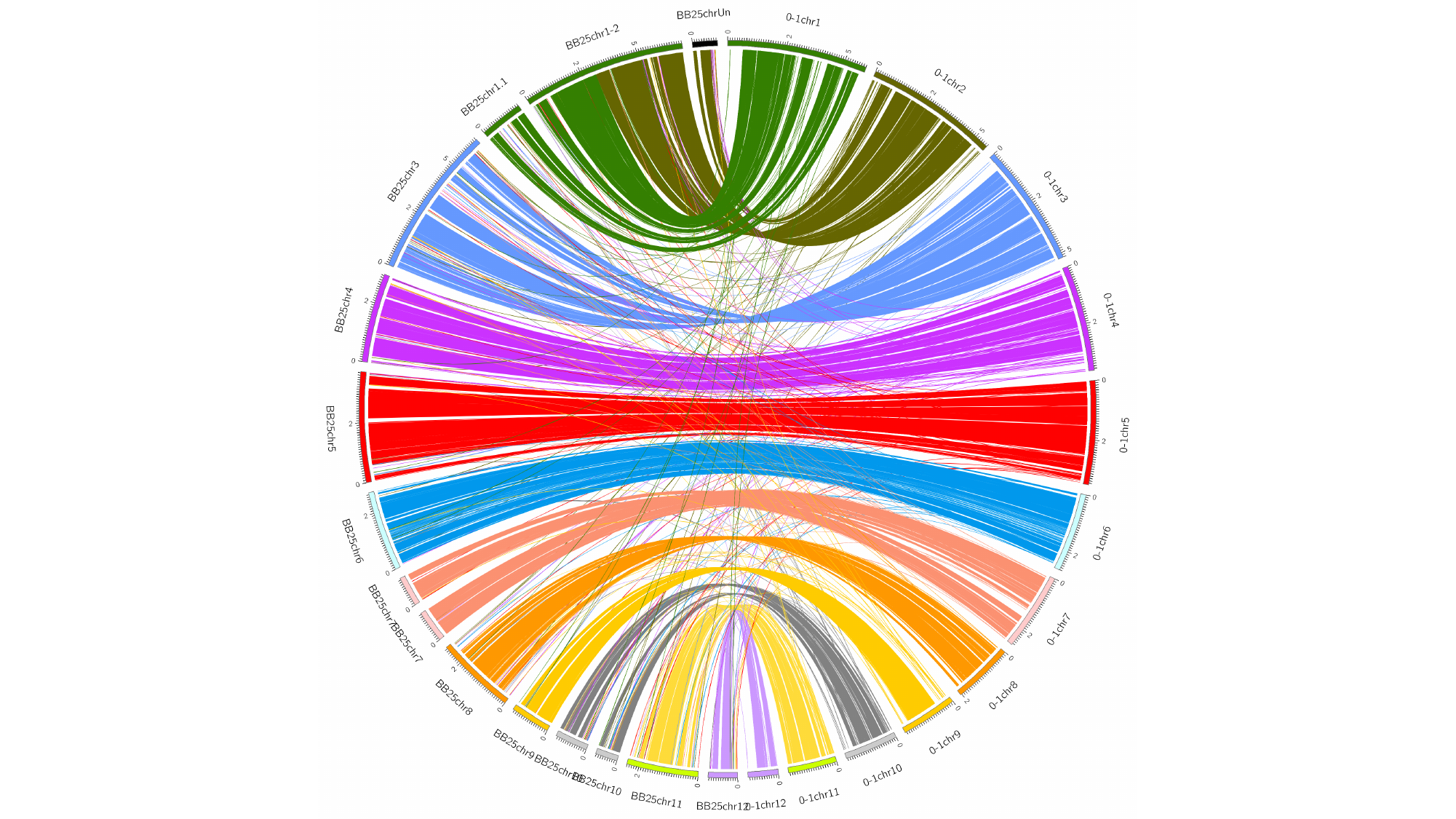
